# Core splicing architecture and early spliceosomal recognition determine microexon sensitivity to SRRM3/4

**DOI:** 10.1101/2024.09.17.613571

**Authors:** Sophie Bonnal, Simon Bajew, Rosa Martinez-Corral, Manuel Irimia

## Abstract

Microexons are essential for proper functioning of neurons and pancreatic endocrine cells, where their inclusion depends on the splicing factors SRRM3/4. However, in pancreatic cells, lower expression of these regulators limits inclusion to only the most sensitive subset among all neuronal microexons. Although various *cis*-acting elements can contribute to microexon regulation, how they determine this differential dose response and high or low sensitivity to SRRM3/4 remains unknown. Here, Massively Parallel Splicing Assays probing 28,535 variants show that sensitivity to SRRM4 is conserved across vertebrates and support a regulatory model whereby high or low microexon sensitivity is largely determined by an interplay between core splicing architecture and length constraints. This conclusion is further supported by distinct spliceosome activities in the absence of SRRM3/4 and by a mathematical model that assumes that the two types of microexons differ only in their efficiency to recruit early spliceosomal components.

## INTRODUCTION

Tissue specific alternative splicing plays a central role in transcriptome and proteome specialization. A set of RNA binding proteins exhibits tissue specific expression patterns and regulates diverse programs of alternative splicing that are instrumental for the development and functioning of specific cell and tissue types^1–3^. One prominent example is that of the paralogous proteins SRRM3 and SRRM4, which are highly expressed in neurons and, in the case of SRRM3, also but to a lesser extent in pancreatic endocrine cells^4^. Both proteins trigger the tissue-specific and dose-dependent inclusion of a highly conserved class of short exons, known as microexons, ranging from 3 to 27 nucleotides (nts)^4–6^. By expressing increasing levels of either SRRM3 or SRRM4, we have previously identified two types of microexons based on their sensitivity to SRRM3/4 regulation. Interestingly, because of their different sensitivity, the low expression of SRRM3 in pancreatic islets is sufficient to drive the inclusion of a subset of highly sensitive (HS) microexons in that tissue, while a higher expression is required to trigger the inclusion of the lowly sensitive (LS), neuronal-specific microexons^4,7^. Despite the functional relevance of the differential sensitivity of microexons to SRRM3/4, the *cis*-regulatory determinants of this sensitivity have not been investigated.

Splice sites are core splicing elements involved in recruiting the spliceosome in a stepwise and dynamic manner. In the early stages of spliceosome assembly, specifically during E complex formation, the 5′ splice site (5′ss) recruits the U1 snRNP through base-pairing interactions between the 5′ end of the U1 snRNA and the 5′ss. Simultaneously, the 3′ splice site (3′ss) mediates the recruitment of the heterodimer U2AF (composed of U2AF1 and U2AF2, associated to the dinucleotide AG and polypyrimidine tract (Py), respectively) and SF1, bound to the branch point (BP) sequence, another key core splicing element. The U2 snRNP is subsequently recruited to the BP through base-pairing interactions between the U2 snRNA and the BP, resulting in the formation of the A complex. Later in the process, the tri-snRNP (U4/U6/U5) is recruited. Multiple RNA-RNA, RNA-protein and protein-protein interactions undergo dynamic rearrangements leading to the formation of the catalytic spliceosome, which ultimately orchestrates the removal of introns and the ligation of exons^8^.

Compared to other exons, SRRM3/4-dependent microexons exhibit distinct features in relation to their adjacent splice sites. They have stronger 5′ss, i.e., more complementary to the 5′ end of U1 snRNA and more similar to the species’ consensus sequence. They also have stronger Py tract and BP signals, but these are located farther away from the AG dinucleotide, which results in a relatively weak 3′ss context^9,10^. This provides space for the accommodation of an intronic splicing enhancer (ISE) composed of a bipartite motif UCU_CUC[N1-50]_UGC capable of binding either positive or negative *trans*-acting regulators of microexon splicing, including SRRM3/4 (associated to UGC), SRSF11, U2AF2 and PTBP1, among others^7,9,11–17^. The contribution of those *cis*-regulatory elements to SRRM3/4-mediated regulation has only been investigated in detail for a few microexons^9,16,18,19^, and never in the context of their differential sensitivity to SRRM3/4.

At least two contrasting yet non-mutually exclusive models taking into account core and SRRM3/4-specific splicing signals may explain the differences in SRRM3/4 sensitivity (Fig. 1a). On the one hand, both classes of microexons (LS and HS) could have similar core splicing architectures (i.e., the organization and strength of their core *cis*-acting splicing elements) but differ in their ability and/or affinity to recruit SRRM3/4 to the pre-mRNA (Fig. 1a, Model I). This model would predict differences in the contribution of SRRM3/4-related features between the two classes of microexons (e.g., UGC presence and/or location). On the other hand, both microexon classes could differ in the composition and strength of core splicing motifs involved in scaffolding the spliceosome, while the recruitment and effect of SRRM3/4 could be similar (Fig. 1a, Model II). This model would predict different effects on each type of microexons if the core splicing architecture is modified in the same manner.

**Fig. 1:**
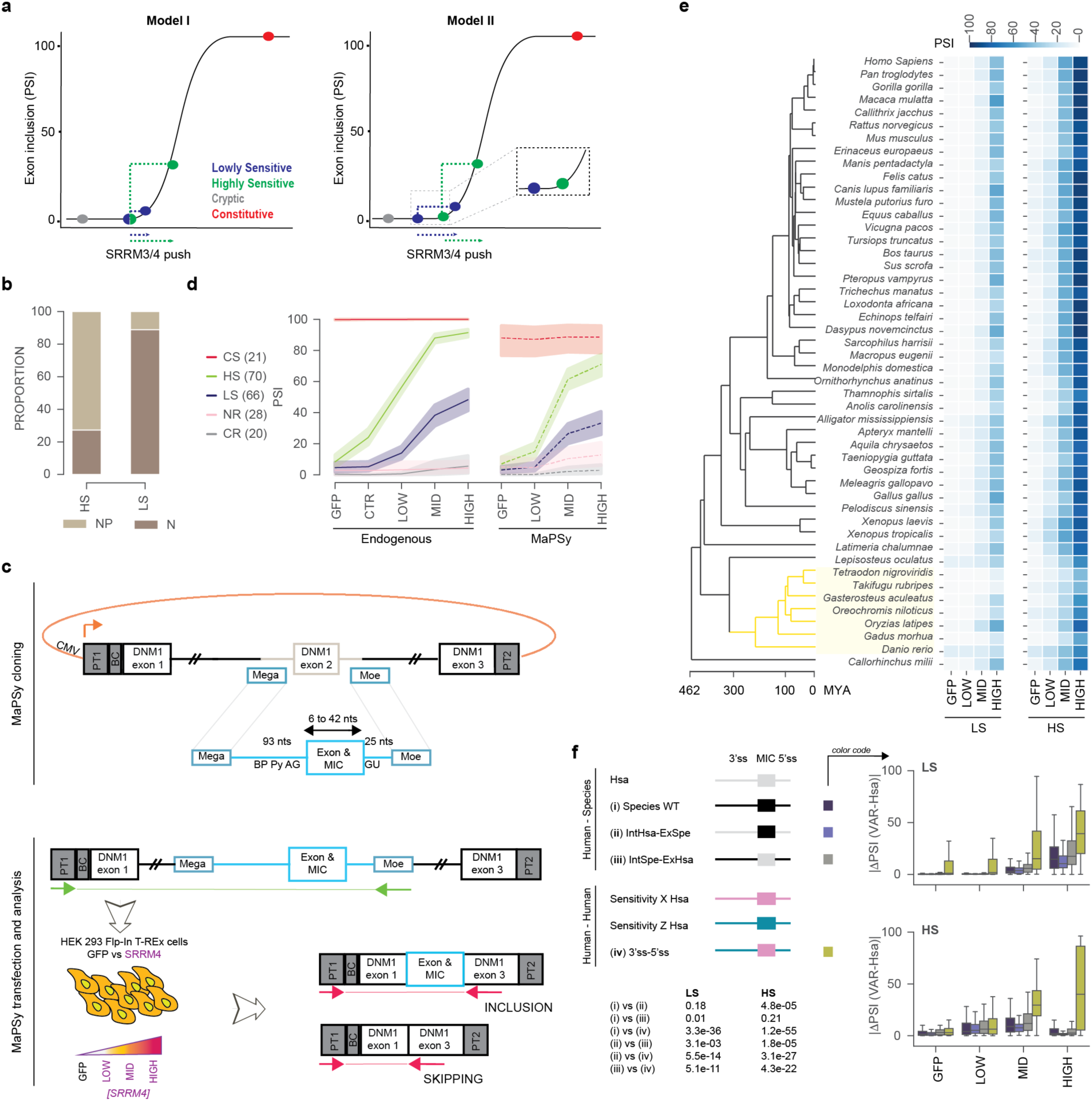
MaPSy strategy recapitulates endogenous microexon sensitivity to SRRM3/4. (**a**) Proposed models of dose-dependent response of microexons with different sensitivities to SRRM3/4. For each exon type, its inclusion level (PSI) in the absence of SRRM3/4 is represented in a PSI continuum, from 0 to 100, which follows a sigmoid curve, as described by^25^. The effect of having a given amount of SRRM3/4 is represented for LS and HS microexons as an enhancing effect, or push, towards higher PSI levels. The difference between the two models resides on the magnitude of this effect and the starting position within this curve for the two types of microexons. According to Model I, the starting position is similar, but the enhancing effect is stronger for HS microexons. In contrast, according to Model II, the effect is similar for both types but HS microexons are shifted towards the right in the PSI curve (as emphasized in the inset), i.e., they are more primed for recognition and inclusion under basal conditions. (**b**) Proportion of LS and HS microexons previously reported as neuronal-specific (N) or neuronal + pancreatic endocrine-specific (NP)^4^. HS = 40 NP + 15 N, LS = 5 NP + 40 N. The remaining microexons were not assigned to either category. (**c**) Overview of the MaPSy strategy for investigating the patterns of microexon splicing *ex vivo*. Pre-designed variants were cloned into a barcoded minigene reporter (top panel) and transfected into HEK 293 Flp-In T-REx cells expressing either GFP (black) or SRRM4 at different levels upon doxycycline titration (purple) (bottom panel). Primers used for sequencing are depicted in green and red (Methods). (**d**) Inclusion levels upon SRRM4 titration in the endogenous (left) or in the MaPSy (right) context for the subset of CS, HS, LS, NR (Non Responding), CR events presented in Supplementary Fig. 1a with sufficient read coverage across all nine experimental conditions (x-axis, Methods). (**e**) The heatmaps represent the mean of the PSI for orthologous WT sequences in four experimental conditions (x-axis; GFP and LOW, MID, HIGH expression of SRRM4) for LS (left) or HS (right) events across a phylogeny of 47 jawed vertebrates (y-axis). Time scale in million years ago (MYA) from TimeTree^64^. The teleost branch is marked in yellow. (**f**) Left: representation of the different intron swappings performed. Variants include: (i) WT orthologous non-human sequences (Species WT), (ii) swapping both flanking intronic sequences from human (Hsa) to another species, (iii) swapping both flanking intronic sequences from another species to human, and (iv) swapping both flanking intronic sequences from one sensitivity group X (LS or HS) to a different sensitivity group Z (HS or LS). The color codes used in the boxplots on the right side are indicated. Right: the boxplots show the absolute change in PSI of each variant type respect to the WT human sequence (|ΔPSI (VAR-Hsa)|). Teleosts were removed from the analysis. Results of Mann-Whitney-Wilcoxon test in HIGH expression level of SRRM4 are indicated for LS and HS events for the indicated comparisons (bottom-left panel). See also Supplementary Fig. 1.

In this study, we have used Massively Parallel Splicing assays (MaPSy) to test predictions made by these two models by assessing the impact of thousands of mutations on microexon sensitivity to SRRM3/4. This type of libraries, consisting of pools of sequences with either random mutations or rationally designed variants, have previously been used by other groups to investigate splicing regulation. These studies focused on investigating the role of *cis*-acting elements in 3′ss selection and regulation upon splicing factor mutation^20,21^, in 5′ss selection and regulation upon perturbations^21–23^, as well as of exonic ^24–37^ or exonic/intronic^38–42^ sequences. Here, we generated and analyzed 28,535 unique variants to assess the evolutionary conservation and regulatory contribution of both intronic (3′ss / 5′ss) and exonic *cis*-acting elements to SRRM3/4 regulation and sensitivity. We report the results of the quantitative dose response of those variants to different levels of SRRM4 activity by titrating its expression in HEK 293 Flp-In T-REx cells. While LS and HS microexons may also exhibit individual differences in their response to changes in SRRM3/4-related motifs, our results are overall highly consistent with the regulatory Model II (Fig. 1a), whereby the main determinants of microexon sensitivity are related to differences in core splicing architecture. This conclusion is further supported by a mathematical model that assumes that the two types of microexons differ in their efficiency of recruitment of early spliceosomal components in the absence of SRRM3/4.

## RESULTS

### MaPSy applied to tissue specific microexons recapitulates SRRM4 splicing regulation

To categorize microexons based on their response to SRRM3/4, we utilized RNA sequencing (RNA-seq) data from two human cell lines (HeLa and HEK 293) expressing either SRRM3 or SRRM4 at four levels and their corresponding control cells. Using Percent Splice-In (PSI) thresholds at each expression level (see Methods), we defined 71 LS and 73 HS SRRM3/4-dependent microexons (Supplementary Fig. 1a, Supplementary Table 1). These microexons were respectively associated with either the neuronal-specific (N) or shared neuronal and pancreatic endocrine cell (NP) specific programs reported previously^4^ (Fig. 1b).

We next developed a MaPSy approach in HEK 293 Flp-In T-REx cell lines to investigate the determinants of the differential sensitivity of LS and HS microexons to SRRM4. Since the main *cis*-acting elements involved in microexon regulation are expected to be upstream and proximal to the dinucleotide AG and at the 5′ ss^9,10,16,18,19^, we designed and cloned each sequence to contain the microexon flanked by the adjacent intronic 93 nts of the 3′ ss and 25 nts of the 5′ ss (Fig. 1c). We then assessed how each class of microexons responded to different levels of SRRM4 expression in the context of the MaPSy libraries. For comparison, we also included 28 non-SRRM3/4-responding (NR) microexons, 20 cryptic (CR) microexons, and 21 constitutively spliced (CS) 42-nt long exons (Supplementary Fig. 1a, Supplementary Table 1; see Methods). The inclusion profiles of exons with sufficient read coverage across all conditions showed that each (micro)exon group exhibited the expected relative behavior compared to their endogenous counterparts, despite an average lower inclusion level across all exon groups (Fig. 1d).

Next, to generate the subsequent variant libraries, we selected a subset of events that overall faithfully recapitulated the endogenous response by taking into account several factors: (i) similarity of the magnitude of response in the minigene with respect to the endogenous, (ii) response of the minigene in the library in both LOW and HIGH conditions, and (iii) low basal inclusion in the minigene in the GFP condition (see Methods), assuming that these constructs carried all the essential *cis*-regulatory elements required for endogenous SRRM4 regulation. A few microexons of known biological interest were also prioritized. We selected 15 HS, 15 LS, 3 CR microexons and 9 CS exons (Supplementary Fig. 1b, Supplementary Table 2), and used them to conduct a targeted high throughput mutagenesis within intronic and exonic sequences. The wild type sequences (WT) and their resulting variants (VAR) were then transfected in HEK 293 cells expressing four levels (none [=GFP], LOW, MID and HIGH) of SRRM4 expression, and RNA-seq was used to quantify the effects of each sequence variant on inclusion levels (Fig. 1c; see Methods).

### Sequence determinants of SRRM4 sensitivity are widely conserved across vertebrates

We first investigated the evolutionary conservation of the *cis*-regulatory architecture underlying the sensitivity to SRRM4. For this purpose, we retrieved the sequences of LS and HS microexons and surrounding splice sites from 47 species covering the main groups of jawed vertebrates across ∼450 million years of evolution (Supplementary Table 3) and generated equivalent minigenes to those of the human counterparts. In line with their high intronic and exonic sequence conservation^5^, relative PSI levels were remarkably conserved from shark to mammals for both LS and HS microexons under different SRRM4 conditions in human cells (Fig. 1e, Supplementary Fig. 1c). Teleosts showed more dissimilar inclusion patterns relative to their corresponding human sequences (Supplementary Fig. 1c), even though their dose response was also conserved. Consistent with this *cis*-regulatory conservation, endogenous orthologous HS microexons exhibited significantly higher overall inclusion levels in neural and endocrine pancreas than LS ones in five studied vertebrates (Supplementary Fig. 1d). Next, to directly test the conservation of the *cis*-regulatory elements in their intronic sequences, we swapped both flanking intronic sequences between each species variant and its human ortholog (Fig. 1f, swap (iii)), or the human event and its ortholog from the other species (Fig. 1f, swap (ii)). As a control, we swapped all flanking intronic sequences between human microexons with different sensitivity (Fig. 1f, swap (iv)). We observed relatively little variation in PSI across those chimeric evolutionary variants compared to the WT human minigene, and this variation was much lower in all cases and conditions compared to intronic swaps among non-homologous human microexons (Fig. 1f, Supplementary Fig. 1e). These results showed that the *cis*-regulatory elements involved in splicing regulation by - and sensitivity to - SRRM4 are largely evolutionarily conserved and capable of recapitulating the expected sensitivity when introduced into human cells.

### Exon length contributes to exon recognition and sensitivity to SRRM4

To start investigating the contribution of various features to SRRM4 sensitivity, and given that microexons are characterized by their short length, we first assessed the impact of increasing or decreasing their length. To increase it, we designed sixteen 36-nt long sequences devoid of potential splicing regulatory motifs such as Exonic Splicing Enhancers (ESE) or Silencers (ESS) according to either the Human Splicing Finder tool^43^ or data from^44^ (“designed sequences”; Methods). Then, for each microexon, we increased the length of the microexon through the stepwise addition of 3 nts to its center for each of the designed sequences, up to a maximal exon length of 42 nts (Supplementary Fig. 2a,b, Supplementary Table 4). As expected, when added to LS, HS and CR events, those sequences led to minimal variation of Exonic Splicing Regulatory sequences (ESRseq) scores throughout those exons^44^ (Supplementary Fig. 2c-f). In parallel, we also decreased the length of each microexon and CS exon by stepwise removal of 3 nts from the middle of the exon until its length reached 6 nts.

This analysis revealed that lengthening generally enhanced exon recognition, as evidenced by the increase in PSI of the longer variants with respect to the WT ones observed across all conditions (Fig. 2a,b; *rho ≥* 0.84 *P* < 0.0001 for all conditions, Spearman Correlation), and despite the unexpected effects of some added sequences (Supplementary Fig. 2g). Conversely, shortening the length had a negative impact on the degree to which the exons responded to SRRM4, as evidenced under conditions of HIGH expression of SRRM4 (Fig. 2a,b). Therefore, this analysis uncovered an overall positive association between length and microexon inclusion in line with previous work^45^. However, we observed that, while both LS and HS events have PSI ∼ 0 in the absence of SRRM4 (GFP condition) (insets in Fig. 2a,b), only HS events showed a strong PSI increase upon lengthening in that condition (Fig. 2b). This indicates that HS microexons can be included in the absence of SRRM4 simply by lengthening, unlike most of LS microexons, whose variants still require SRRM4 to achieve a substantial inclusion level.

**Fig. 2:**
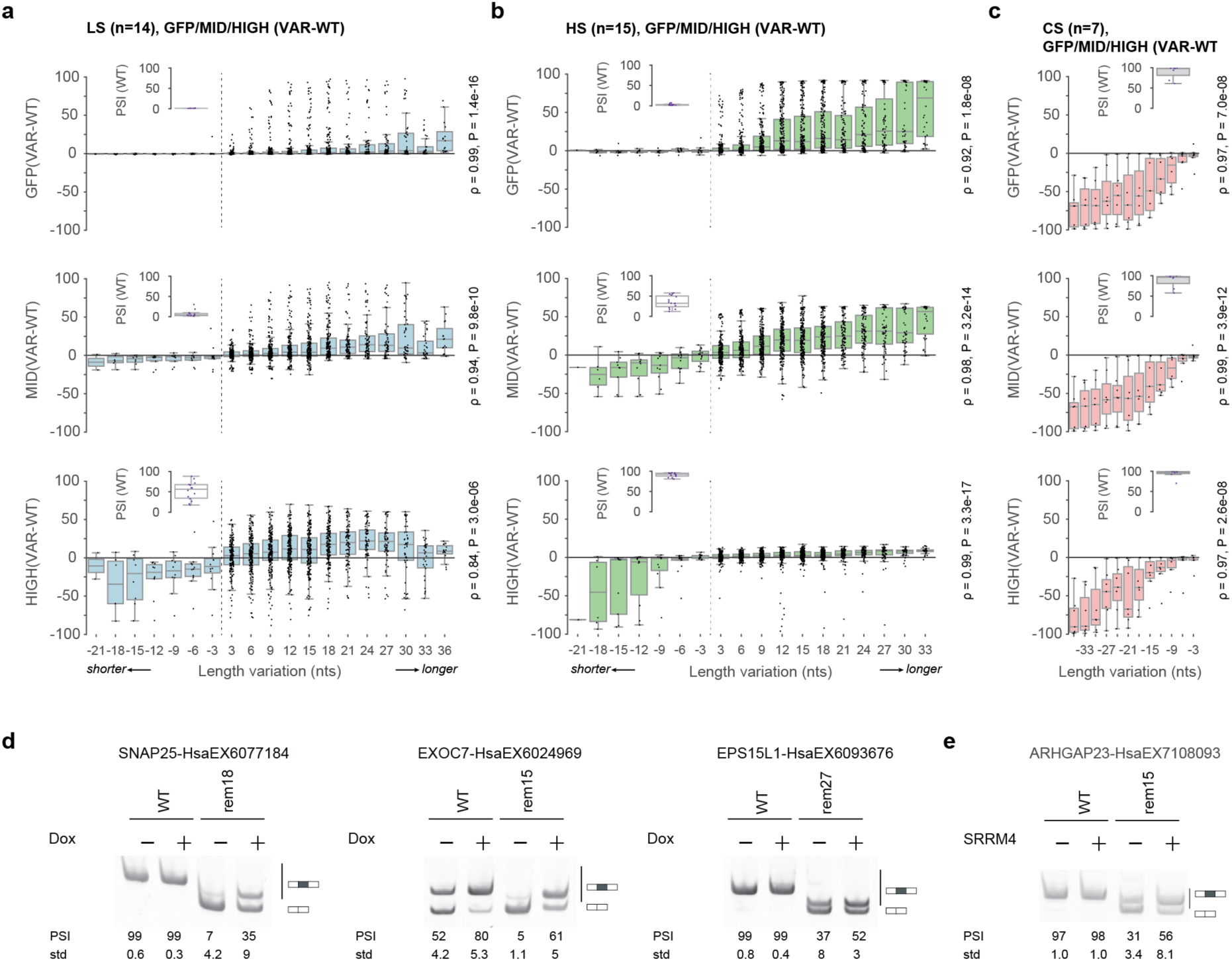
Exonic length contributes to splicing regulation in response to SRRM3/4. (**a-c**) Impact of exon length on the pattern of splicing and regulation by SRRM4 for LS (a), HS (b) and CS (c) events. Change in exon inclusion upon shortening or lengthening was quantified as ΔPSI (VAR-WT) (i.e., difference in PSI between the different length VARiants and their WT counterpart) in either the GFP condition or under MID, HIGH level of expression of SRRM4. Each data point corresponds to the effect of adding a given number of nucleotides from one of the fifteen designed sequences (positive number on the x-axis), or removal of a given number of nucleotides from the exonic sequence (negative number on the x-axis). The boxplots in the insets represent the distribution of PSIs of the WT constructs in each experimental condition. In (a,b), the reference length of the WT is indicated by the dashed vertical line. (**d,e**) RT-PCR assays showing the splicing pattern under control condition (-) or expression of SRRM4 (+) of four CS events (WT length of 42 nt) and their length variants, in which the indicated number of nucleotides, N, has been removed (remN). PSI and standard deviations (std) from five biological replicates are provided. In (d), HEK 293 Flp-In T-REx cells non treated or treated with doxycycline were used. In (e), transient expression of a control vector (-) or SRRM4 expression vector (+) in HEK 293 cells was performed. See also Supplementary Fig. 2,3.

We next examined the consequence of reducing the size of the 42 nt-long CS exons. In general, all studied events displayed progressively reduced inclusion under the GFP condition as the length decreased (Fig. 2c). However, the specific length at which inclusion started to decrease and when it reached nearly zero varied for each event (Supplementary Fig. 2h,i). Strikingly, we also noticed that some of the CS events became responsive to SRRM4 at a certain length (ΔPSI (HIGH-GFP) ≥ 20; Supplementary Fig. 2h). The gain of SRRM4 regulation by shortened CS variants was independently validated for four of these events through individual transfection in HEK 293 cells expressing either GFP or SRRM4 (Fig. 2d,e and Supplementary Fig. 2j). Finally, CR microexons also exhibited a positive response to lengthening while the shortening had a more dispersed pattern (Supplementary Fig. 2k,l).

Altogether, these results show that microexon length is one of the determinants of exon recognition and contributes to sensitivity to SRRM4, and it does so differently for HS and LS microexons. For HS microexons, the sensitivity is largely maximized while increases in length often lead to substantial microexon variant inclusion even in the absence of SRRM4. For LS microexons, longer variants have increased sensitivity to SRRM4, but they do not usually achieve high inclusion levels in the GFP condition.

### Exonic elements contribute to microexon regulation in a context dependent manner

*Cis*-acting elements located within exons are known to contribute to splicing regulation^29,33,44^. As we observed distinct effects for some sequences used for microexons lengthening (Supplementary Fig. 2g), we next asked whether exon-specific regulatory motifs were present. For that purpose, we performed a systematic mutagenesis of three successive nucleotides using a sliding window approach from the 5′ end to the 3′ end of each microexon (Supplementary Fig. 3a). Interestingly, similar to the pattern observed for lengthening, only a few mutations in HS microexons showed substantial positive responses (ΔPSI > 10) in the absence of SRRM4 (Supplementary Fig. 3b,c,e, *P* = 1.2e-4 two-sided Fisher’s Exact test). Upon different levels of SRRM4 expression, mutations with strong effects were still a minority, but these were balanced between those with a positive and a negative effect on SRRM4 regulation for both LS and HS microexons (Supplementary Fig. 3b,c,e,f). In the case of CS exons and their corresponding shorter variants, we noticed exon length-dependent effects for some *cis*-acting elements, which only contributed substantially to splicing regulation in the shortened variants (Supplementary Fig. 3d,e). Moreover, we also detected specific mutations that enabled regulation by SRRM4, even for the 42 nt-long exon variants (Supplementary Fig. 3f).

To complement this mutational scan, we created another set of variants by inserting previously described ESE or ESS hexamers^44,46,47^ at two exonic positions (Supplementary Fig. 3g). As expected, ESE and ESS insertions usually led to increased and decreased (micro)exon inclusion, respectively, with some context dependent effects observed across all exon types and conditions. However, we again observed that ESE insertion led to substantial SRRM4-independent inclusion only for HS but not LS microexons (Supplementary Fig. 3g). Interestingly, this pattern was also observed for shortened SRRM4-dependent CS events in the GFP condition (Supplementary Fig. 3g).

Altogether, these results indicate that, while it is likely that microexon regulation mainly relies on elements in the flanking intronic regions, *cis*-regulatory elements within the microexon may also impact its inclusion and response to SRRM4.

### The sensitivity to SRRM4 is to a large extent encoded in the flanking intronic sequences

To assess to what extent the different sensitivities to SRRM4 are encoded in the flanking intronic sequences, we systematically swapped the 3′, the 5′ or both flanking intronic sequences among all 15 HS and 15 LS studied microexons (3,480 total combinations) (Fig. 3a). These swappings showed that sensitivity information was encoded in both the 3′ and, less expectedly, the 5′ neighboring intronic sequences (Fig. 3). Replacing the intronic sequences from HS events with those from LS events negatively impacted the response to SRRM4 (Fig. 3b), while the opposite was observed when replacing LS with HS intronic sequences (Fig. 3c). Surprisingly, the effect was more pronounced when swapping the 5′ss as compared to the 3′ss and even more pronounced when both flanking intronic regions were replaced at given concentrations of SRRM4 (Fig. 3a-c). These results thus point towards a more important role of the 5′ss than previously anticipated^5,9,16^. Swapping sequences within HS events or within LS events yielded more moderate impacts (Fig. 3a-c), suggesting the presence of more compatible intronic architectures for a given sensitivity.

**Fig. 3:**
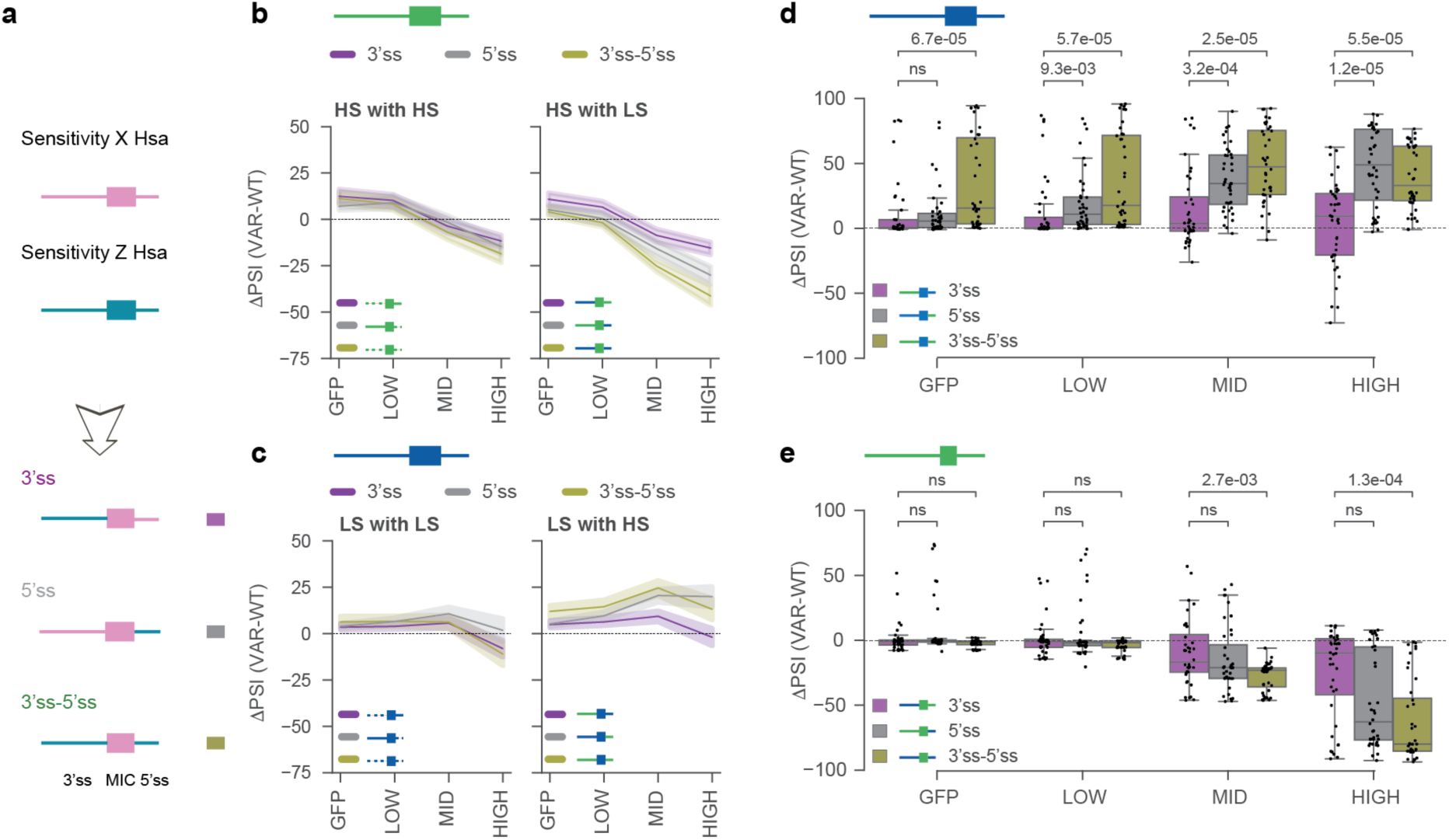
Intronic elements contribute to the different microexon sensitivities. (**a**) Representation of the different intron swappings performed in (b-e). Flanking introns from events in Sensitivity group Z are used to replace the corresponding regions in events from Sensitivity group X. Color codes used in (b-e) are indicated. (**b-e**) Effect of swappings intronic sequences between events from Sensitivity group X and Sensitivity group Z. (b) group X: HS, group Z: HS, LS. (c) group X: LS, group Z: LS, HS. (d) group X: LS long (i.e., exon length > 15 nt), group Z: HS short (i.e., exon length < 16 nt). (e) group X: HS short, group Z: LS long. LS and HS pre-mRNA regions are represented in blue, and green, respectively. For each subpanel, the WT sequence is depicted at the top and a scheme of the part(s) of the pre-mRNA involved in the swapping is represented along with the color legend of the lines or boxes. Statistics in (b,c): results of one sample t-tests are presented in Supplementary Table 6, in (d,e): Mann-Whitney-Wilcoxon test two-sided with Bonferroni correction (ns: not significant).

Given the importance of length on microexon inclusion (Fig. 2), we next focused on swappings between events of comparable length (both ≤ 15 nts, short; or both > 15 nts, long) or between events of different length groups (short with long and vice versa). Remarkably, this analysis revealed that the *cis*-regulatory architecture of HS and LS is adapted to the length context of the microexons. Therefore, when HS intronic sequences from short microexons (predicted to be the most sensitive) were placed around long LS microexons (predicted to be the least sensitive), we observed an overall increase in PSI across all conditions, including a substantial inclusion even in the absence of SRRM4 (Fig. 3d). On the contrary, when LS intronic sequences from long microexons were placed around short HS microexons, we found an overall decrease in inclusion, particularly noticeable under HIGH SRRM4 levels (Fig. 3e). However, the effect of replacing the 5′ss was mainly bimodal in the HIGH condition: while 37.5% of variants had a strong effect (ΔPSI < −75), 12% had nearly no change upon swap (ΔPSI > −10).

These experiments show that the sensitivity to SRRM4 is to a large extent encoded in the intronic sequences surrounding the microexons, which prompted us to investigate the relevance of specific *cis*-acting elements globally or individually within these intronic sequences.

### Context and position requirements for the UGC motif are similar between sensitivity groups

Next, we focused on the other highly characteristic feature of SRRM3/4-regulated microexons, the UGC motif near the 3′ss. A UGC motif between the BP and the AG was found in the majority, but not all, SRRM3/4-dependent microexons. However, its presence is not associated with their sensitivity, since a UGC motif was found in 79% (57/73) HS and 74% (51/71) LS microexons (*P* = 0.444 two-sided Fisher’s Exact test). To systematically explore the role of this motif in SRRM3/4-dependent splicing regulation, we mutated either one or multiple UGCs within the upstream intronic sequences of each human microexon used in our mutagenesis study. Consistent with previous reports, most single or multiple mutations of UGC motifs disrupted the response to SRRM4 for both LS and HS events (Fig. 4a). Moreover, when multiple UGC motifs were present, not all of them had an impact when disrupted (Fig. 4a).

**Fig. 4:**
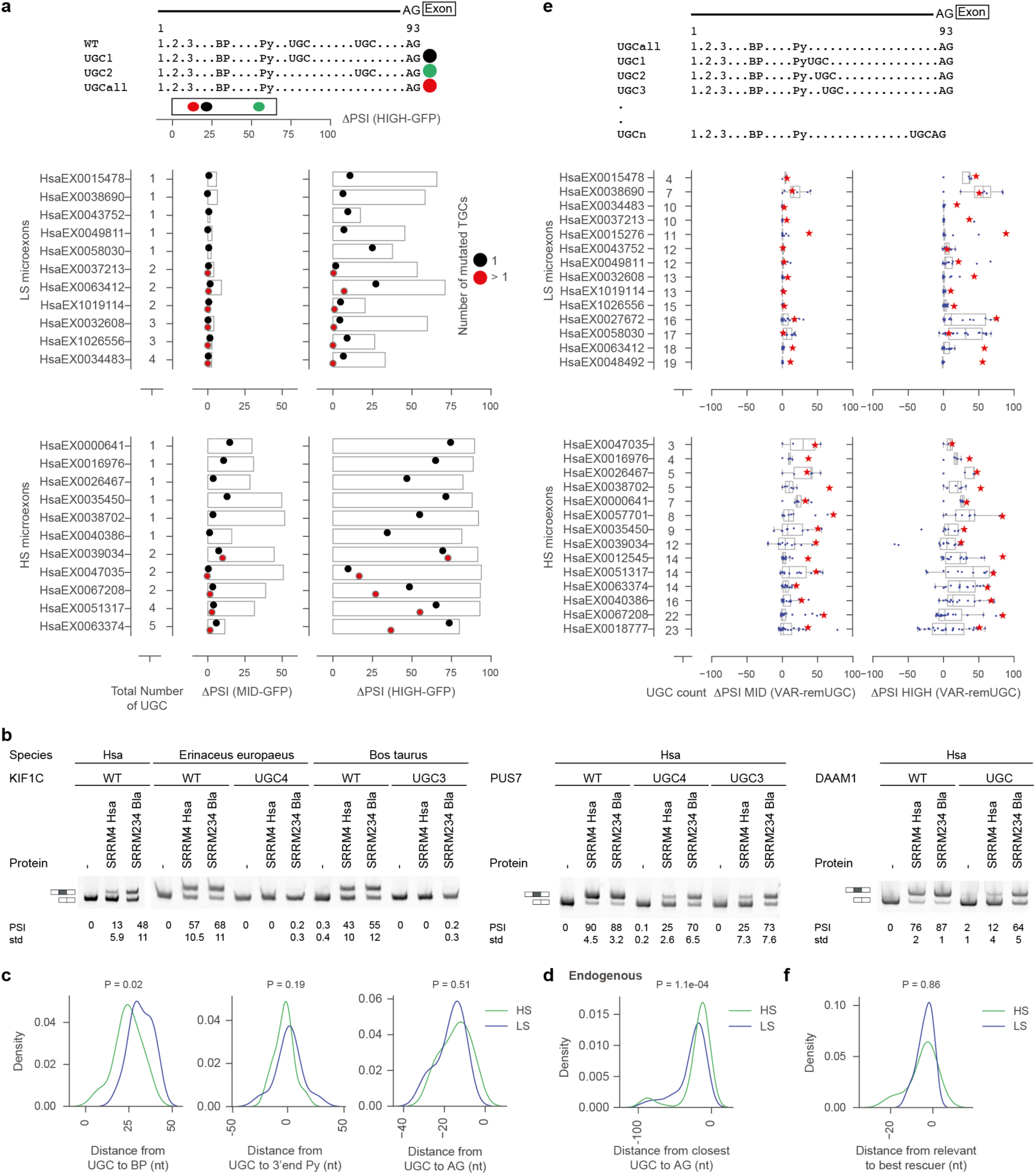
The UGC motif contributes to splicing regulation by SRRM4 in a context and position dependent manner. (**a**) Top: Scheme of the generation and quantification of UGC variants. UGC motifs in the 3’ss of the WT events were mutated either individually (black and green dots) or in combinations (red dot). Bottom: Quantification of the impact of UGC mutations for LS and HS microexons. The barplots represent the ΔPSI of the WT sequences in MID (left) and HIGH (right) expression of SRRM4 with respect to GFP. The effect of mutating either the UGC with the strongest impact (black dots) or all UGCs (red dots, total number considered for mutagenesis is provided as “Total number of UGCs”) is shown as the ΔPSI between MID or HIGH expression of SRRM4 with respect to GFP for each variant. The closer to x = 0, the stronger the effect of the UGC mutation(s). (**b**) RT-PCR assays showing the splicing pattern of three events (WT and UGC variant) from different species under control condition (-) or expression of human SRRM4 (Hsa) or amphioxus SRRM234 (Bla). PSI and standard deviations (std) from three biological replicates are provided. The PSI response of KIF1C-HsaEX0034483 and PUS7-HsaEX0051317 variants in MaPSy is presented in Supplementary Fig. 4b. (**c**) Distribution of the distances between the most relevant UGC motif (as identified in (a), black dot) and the dinucleotide AG, the 3′ end of the polypyrimidine (Py) tract or the best predicted branch point (BP). The UGC-AG distance is expressed in negative values to account for the reverse arrangement of the two elements with respect to BP-UGC and Py-UGC pairs. (**d**) Distribution of the distances between the dinucleotide AG and the closest UGC for all endogenous LS and HS events with UGC in the 93-nt long flanking 3′ss. (**e**) The UGC-addition walk (top panel) examines the effect of adding a single UGC between the 3′ end of the Py tract and the dinucleotide AG. This effect is quantified as ΔPSI with respect to the variant in which all UGCs are mutated (UGCall in panel (a)) in MID or HIGH expression levels of SRRM4. The red stars represent the WT and the blue dots the variants of the UGC-addition walk, whose distribution is shown as a boxplot. (**f**) Distribution of the distance between the most relevant UGC identified in (a) (black dots) and the UGC with the strongest effect identified in (c). Statistics in (c,d,f): Mann-Whitney-Wilcoxon test. See also Supplementary Fig. 4.

Surprisingly, however, the presence of a UGC motif was not strictly necessary for SRRM4-dependent inclusion of microexons, in contrast to previous reports^6,9,10,18,19^. In particular, HIGH levels of SRRM4 could, to varying extents, overcome the disruption of UGC motifs for most microexons. In such cases, HS microexons displayed higher propensity for inclusion despite the mutation of UGC motifs (Fig. 4a). Moreover, a similar pattern was observed for orthologous microexons from 23 other species, for which all present UGC motifs were mutated (Supplementary Fig. 4a). The patterns of splicing were independently validated by RT-PCR assays for three microexons after co-expression of either human SRRM4 or a more efficient regulator obtained from amphioxus^10^ (*Srrm234* isoform AB) (Fig. 4b). These results suggest that, while the UGC motif seems to strongly facilitate the action of SRRM3/4, likely favoring its binding, it is not strictly necessary for it. This points towards the presence of additional interactions triggering its recruitment to microexons.

Next, we assessed the relative position of the most relevant UGC, as identified by mutagenesis (Fig. 4a), with respect to the AG, best predicted BP and 3′ end of the Py tract. We found that the relative position of the UGC motif was similar between LS and HS events (Fig. 4c), although the UGC was closer to the BP in HS microexons, suggesting a more compact architecture. In line with this, when all endogenous microexons were considered, the distance between the closest UGC and the AG was also significantly shorter for HS events (Fig. 4d; *P* = 1.1e-04, Mann-Whitney-Wilcoxon test). To assess whether the most relevant UGC was located at the most optimal position in both types of microexons, we performed a “UGC-addition walk”, i.e., we systematically inserted a UGC motif at all possible positions between the 3′ end of the Py tract and the dinucleotide AG after mutating all natural UGC motifs, and assessed the inclusion level of the resulting microexon variants (Fig. 4e, Methods). This analysis showed that for both types of microexons, the most relevant natural UGC is generally at the position where a UGC motif has its maximal effect, even if UGCs at neighboring positions can have similarly strong effects (Fig. 4e,f). Importantly, this result also indicates the UGC motifs are not placed in more optimal configurations in HS than in LS microexons with regard to the general architecture of each exon, i.e., their core splicing elements, as potentially predicted by Model I (Fig. 1a).

Interestingly, for some experimentally tested cases in which an alternative UGC position was more optimal than the WT one, we found that they can partly compensate for the decrease in inclusion caused by shortening the microexon (Supplementary Fig. 4c). This supports a combinational effect of the different features determining SRRM4-mediated microexon inclusion. Finally, we also assessed how additional UGC motifs impacted microexon inclusion and the degree of response to SRRM4. For this purpose, we added one, two or three UGCs downstream to the best predicted BP alone or in combinations with upstream shifts of the BP and its associated Py tract (Supplementary Fig. 4d,e and Methods). Somewhat surprisingly, most of these mutations had a detrimental effect on the response to SRRM4 for both HS and LS events (Supplementary Fig. 4d,e), likely because they led to either decreased Py tract strength or increased distance from BP to AG.

Overall, these findings show that a UGC motif at a specific location is of high relevance for SRRM4 response and this location is largely equivalent for HS and LS microexons. However, the presence of this motif is not strictly necessary for the inclusion of either HS or LS microexons. Therefore, additional *cis*-regulatory elements likely contribute to the splicing pattern and the regulation by SRRM4.

### Sequences in proximity to splice sites are key for SRRM4 splicing regulation

To investigate the parts of the pre-mRNA that are most relevant for microexon splicing and their regulation by SRRM4, we performed a semi-deep mutagenesis scan of both intronic and exonic sequences (Fig. 5a, Supplementary Fig. 3a-c). Overall, nearly all mutations had a neutral or negative impact on splicing. As expected, intronic mutations in proximity to the 3′ss (i.e., likely within the BP, Py tract and intronic splicing regulatory elements) or the 5′ss more negatively impacted the regulation of LS and HS events by SRRM4 (Fig. 5a, Supplementary Fig. 5a). Similar trends were observed for CS events and their shortened variants (Supplementary Fig. 5b), although strong negative effects on inclusion were also observed for WT CS exons in the GFP condition, given their high starting PSI. Moreover, impactful mutations at the 3′ss of CS events tended to be located closer to the dinucleotide AG in comparison with SRRM4-dependent microexons, consistent with the peculiar architecture of the latter, with more distant BPs (see Introduction, Supplementary Fig. 5c).

**Fig. 5:**
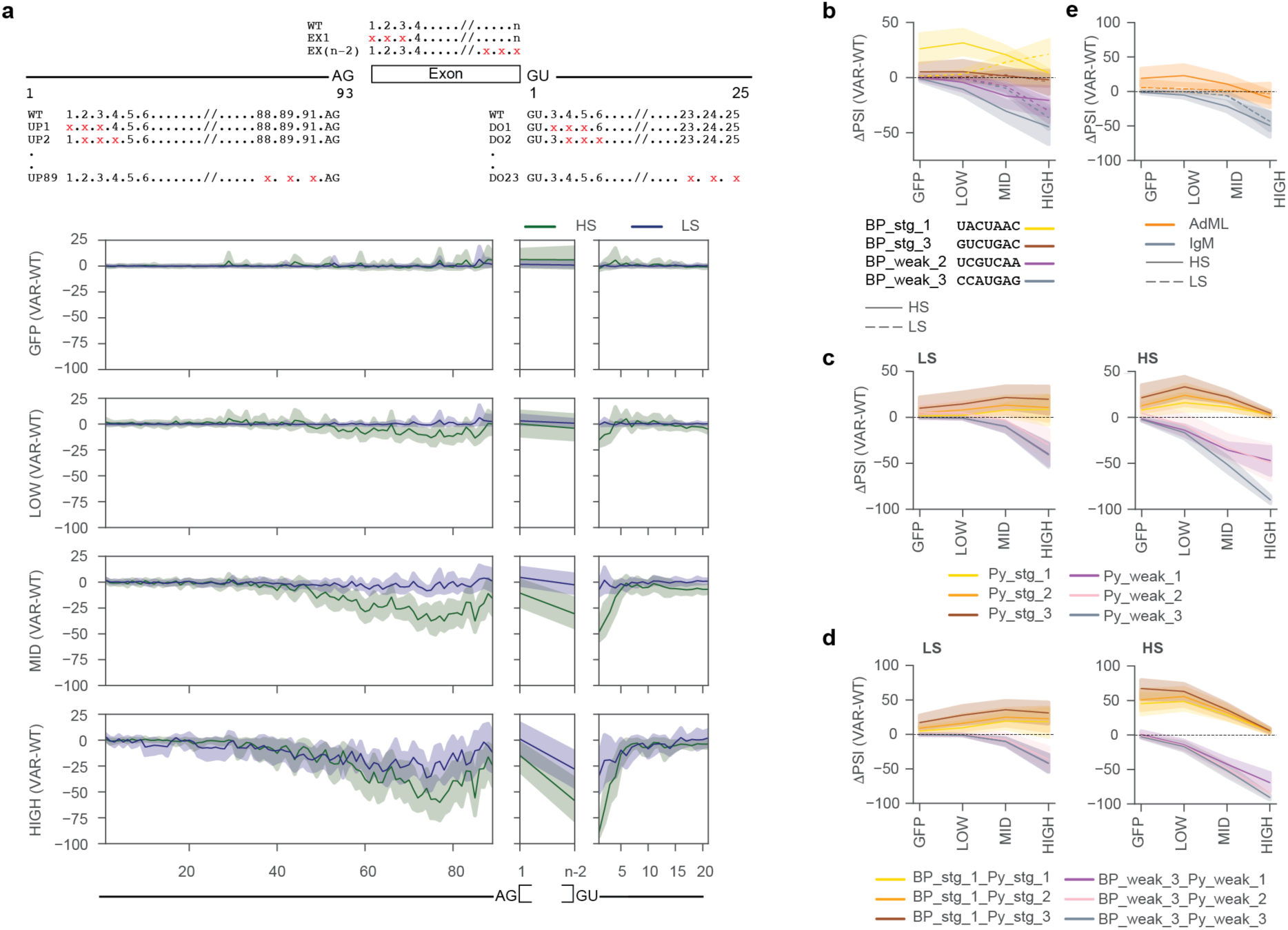
Intronic core splicing elements close to splice sites are relevant for regulation by SRRM4. (**a-e**) Effects of (a) mutating three consecutive nucleotides following the strategy depicted at the top, (b) strengthening (stg) or weakening (weak) the BP (Supplementary Table 7), (c) strengthening (stg) or weakening (weak) the Py tract, (d) combining the mutations described in (b) and (c), or (e) replacing the endogenous BP-Py tract sequence by the ones from AdML or IgM. The effects are shown for LS and HS events for the four experimental conditions quantified as ΔPSI (VAR-WT) (i.e., PSI of the VARiant with respect to the WT sequence). In (a), the positions of the intronic mutations are indicated along the x-axis according to the nomenclature described at the top and only the impact of mutating the first (1) or last 3 (n-2) exonic nucleotides is reported here. The impact of other exonic mutations is reported in Supplementary Fig. 3a-c. See also Supplementary Fig. 5. In (b), the BP sequences used are indicated. For other mutations, see Supplementary Table 5. Statistics in (b-e): results of one sample t-tests are presented in Supplementary Table 6.

To assess the role of the strength of the BP and the Py tract in microexon splicing and regulation by SRRM4, we then replaced the BP of LS and HS events by either two different strong or two weak BPs. As expected, this revealed a general increased or decreased response upon strengthening or weakening, respectively (Fig. 5b). However, the strengthening effect of the most consensus BP (UACUAAC) was particularly remarkable for HS microexons in the GFP condition, and it revealed that modulating core splicing signals is sufficient to generate SRRM4-independent inclusion specifically for HS microexons. In the case of LS microexons, the strengthening of the BP resulted in an overall increase in sensitivity, with very little impact on the PSI in the GFP condition (Fig. 5b). Hence, solely by modulating this core splicing signal, we could convert LS into HS-like microexons, as predicted by Model II (Fig. 1a). Conversely, weakening of the BP signal was sufficient to reduce the sensitivity to SRRM4 of HS and LS microexons (Fig. 5b).

Strengthening or weakening the Py tract^48^ (Supplementary Table 5), alone or in combination with the above mentioned BP mutations, also positively or negatively impacted splicing regulation, respectively. Again, this resulted in SRRM4-independent inclusion, specially for HS microexons, and to the expected changes in sensitivity for both HS and LS microexons (Fig. 5c,d). The joint contribution of the BP and Py tract to regulation by SRRM4 was further confirmed by the full replacement of both elements by the one from Adenovirus Major Late (AdML)^49^ or from immunoglobulin mu (IgM)^50^, two well established splicing substrates with strong or weak 3′ss features, respectively (Fig. 5e). Interestingly, a similar behavior to HS microexons was observed for the SRRM4-dependent microexons obtained from reducing the length of CS events, suggesting that their initial constitutive architecture may be closer to that of HS than of LS microexons (Supplementary Fig. 5d,e). Finally, other probed elements, including UCUC motifs involved in SRSF11 / PTBP1 mediated regulation of microexons^9,16^, did not have a major consistent impact on the tested microexons (Supplementary Fig. 5f).

Altogether, these results show that core elements involved in spliceosomal recruitment contribute to the splicing pattern of microexons and to their sensitivity to SRRM4.

### Unappreciated contribution of the 5′ splice site to splicing regulation by SRRM4

We next assessed more specifically the contribution of the 3′ss and 5′ss sequences. We first focused on the 3′ss, which is recognized by U2AF1 based on positions −3 to +1 (YAG|R^51–53)^. Interestingly, microexons have similar features to AG-independent exons^54^ and have been shown to be generally repressed by U2AF1^13^. Thus, we performed a deep mutagenesis of positions −3, +1, +2 and +3 to globally examine the contribution of the nucleotides surrounding the AG to the regulation by SRRM4. Overall, we noticed microexon-specific dependencies on certain nucleotides (Supplementary Fig. 5g). Also, somewhat unexpectedly, the shortest variants of the CS events were more affected than their longer counterpart, and, to a similar extent as LS and HS events (Supplementary Fig. 5g). These results suggest a complex and microexon-specific relationship between AG recognition and SRRM4-dependent regulation that would require further investigation.

Next, we explored the 5′ss. While it was previously noted that the 5′ss of SRRM3/4-dependent microexons are generally strong^10^, their role in shaping splicing patterns and their contribution to SRRM4 regulation have not been experimentally investigated. As shown above, swaps of intronic regions including the 5′ flanking intronic region had stronger effects than swapping only the upstream intron (Fig. 1f, 3b,c), and mutations in the 5′ss had highly detrimental effects (Fig. 3, 5a). Thus, we sought to determine whether strengthening the 5′ss alone could influence SRRM3/4-dependent splicing regulation. For this purpose, we increased the 5′ss strength by introducing mutations in the last three nucleotides of the exon and up to the eight first nucleotides of the downstream intron, enhancing its complementarity to U1 snRNA (U1cons) (Fig. 6a). Additionally, we performed mutations from positions +5 to +9 in the intron to enhance the complementarity to U6 snRNA (U6cons)^55^ (Supplementary Fig. 6a). Each of those mutations led to a significant increase in the maximal entropy (MAXENT) score of the 5′ss^56^ (Fig. 6b, Supplementary Fig. 6b,c). Strengthening of the 5′ss had a very strong positive effect on inclusion for both types of microexons across most SRRM4 conditions (Fig. 6c,d). In the case of LS microexons, their inclusion reached near saturation at HIGH SRRM4 levels, similar to HS events, and both LS and HS events increase their inclusion levels at low levels of SRRM4, and thus their sensitivity to the regulator. Moreover, we observed a substantial degree of inclusion in the GFP condition for both LS and HS microexons, showing that modulating 5′ss signals is sufficient to drive inclusion of microexons even in the absence of SRRM4. These findings were independently validated by RT-PCR assays for two LS microexons (Fig. 6e, Supplementary Fig. 6d,e). To rule out a potential impact of the specific minigene design on these findings, we validated the impact of strengthening the 5′ss using three additional minigenes for which the endogenous upstream exon/intron, microexon and downstream intron/exon belonged to the same event (Supplementary Fig. 6f).

**Fig. 6:**
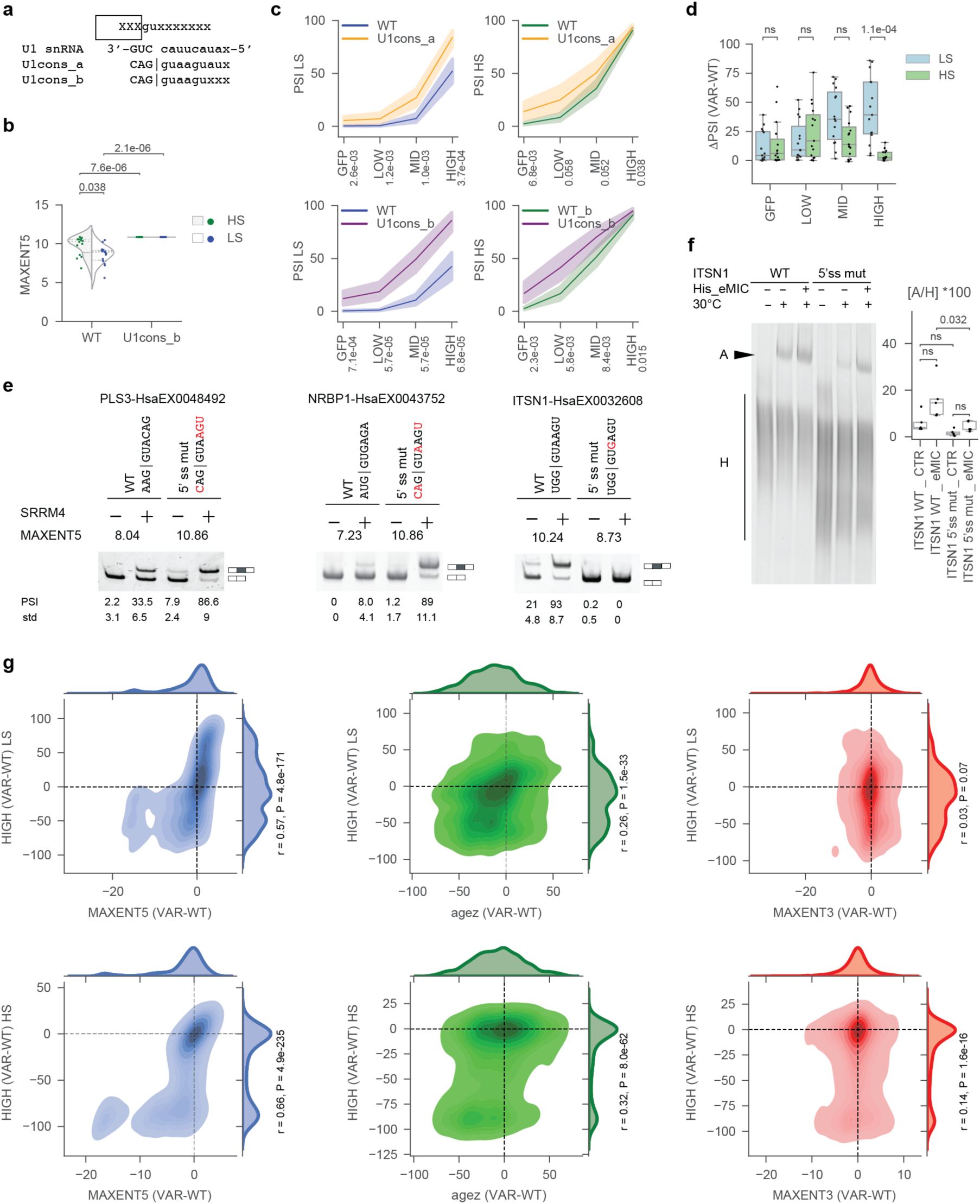
The strength of the 5′ss contributes to sensitivity to SRRM4 expression. (**a**) Mutations performed to enhance base-pairing between the 5′ss and U1 snRNA. U1cons_a corresponds to 9 nts complementarity to U1 snRNA(Yeo and Burge 2004), while U1cons_b corresponds to an extended based pairing involving MAG|GURAGUAU^70^. (**b**) Maximum entropy scores of the 5′ss (MAXENT5) for WT and U1 variants for LS and HS events. (**c**) Splicing profiles quantified as PSI in four experimental conditions for WT and U1 variants depicted in (A) for LS and HS events. Statistics: Mann-Whitney-Wilcoxon test. (**d**) The effects of U1cons_b mutations represented in (a, c) quantified as ΔPSI (VAR-WT) in four experimental conditions for LS and HS events. Each dot represents an event. (**e**) RT-PCR assays showing the splicing pattern of the WT and corresponding U1 variants for three events under control condition (-) or expression of SRRM4 (+). Sequences of the 5′ss are indicated at the top and the mutated nucleotides are highlighted in red. PSI and standard deviations (std) from at least three biological replicates are provided. PSI in the MaPSy are provided in Supplementary Fig. 6d,e. (**f**) *In vitro* spliceosomal A complex formation on ITSN1 pre-mRNA, either WT or the 5′ss variant in (e), under control condition or expression of the recombinant eMIC domain of SRRM3^10^. Quantification of the ratio A/H complexes from five biological replicates is provided (Supplementary Table 8). (**g**) Distribution of ΔPSI (VAR-WT) values upon HIGH level of expression of SRRM4 with respect to ΔMAXENT5 (VAR-WT), Δagez (VAR-WT), ΔMAXENT3 (VAR-WT) values for the LS and HS variants with WT exon length and that modify each corresponding feature (see Methods). MAXENT, maximum entropy score; agez, AG exclusion zone. Statistics in b,d,f: Mann-Whitney-Wilcoxon test two-sided with Bonferroni correction (ns: non significant). Statistics in g: Pearson correlation coefficients and p-values. See also Supplementary Fig. 6.

The reverse pattern was observed upon weakening the strength of the 5′ss by GU to GC mutation or by using a weak 5′ss sequence (Supplementary Fig. 6a,g,h). To exemplify these effects, we focused on WT orthologous sequences for the LS microexon in *ITSN1* (HsaEX0032608). We noticed that the orthologous microexon for pig, cow and dolphin (belonging to the monophyletic group of Artiodactyls) had the lowest response to SRRM4 among twenty probed species (Supplementary Fig. 6i, right panel). A multiple sequence alignment revealed an A-to-G substitution at position +3 of the 5′ss in this clade (Supplementary Fig. 6i, left panel), which decreased its maximal entropy score from 10.24 to 8.73 (Fig. 6e). To test the effect of this variant in the response to SRRM4, we generated an A-to-G single point mutation at position +3 of the human sequence, which led to a complete loss of response to SRRM4 expression (Fig. 6e). The contribution of the 5′ss in early spliceosomal recruitment was further examined *in vitro* for this event. The A complex formation efficiency on *ITSN1* microexon substrates in the presence of the minimal domain of SRRM3 necessary and sufficient for regulation (eMIC) was higher when a stronger 5′ss was present (Fig. 6f), suggesting a potential contribution of exon definition in the regulation. Another set of examples was provided by two likely pathogenic SNPs, according to ClinVar, which weaken the 5′ss and led to decreased regulation of a HS microexon in *CASK* (HsaEX0012545) and a CS exon in *SNAP25* (HsaEX6077184) (Supplementary Fig. 6j). These patterns are consistent with the global results obtained by performing deep mutagenesis of every position involved in base pairing to U1 snRNA (from −3 to +6) (Supplementary Fig. 6k). Interestingly, while mutations at positions −1 and +3 were detrimental for all types of (micro)exons, only HS and shortened CS events were highly sensitive to mutations at position +5 (Supplementary Fig. 6k).

Next, to systematically evaluate the contribution of the 5′ss to the regulation of SRRM4-dependent microexons, we calculated the difference in scores measuring either the 5′ss strength (MAXENT5), the length of the AG exclusion zone (agez) or the 3′ss strength (MAXENT3) between a given variant and its corresponding WT, and correlated it with the change in exon inclusion at HIGH SRRM4 expression between that variant and the corresponding WT. For both LS and HS events, we observed a strong positive correlation only between the variation in MAXENT5 and the change in inclusion, while the correlations with the agez and, specially, MAXENT3 were much weaker (Fig. 6g).

Altogether, these results highlight a previously unappreciated relevance of 5′ss strength in inclusion of the microexon and response to SRRM4 expression.

### Early recognition of splice sites in the absence of SRRM4 correlates with microexon sensitivity

A number of findings presented thus far collectively support a model by which HS and LS microexons differ in their recognition capacities by the spliceosome under conditions where SRRM4 is not expressed. Specifically, HS microexons were more often included in the absence of SRRM4 (GFP condition) upon different types of mutations. This occurred when the microexon length was increased (Fig. 2), an ESE was introduced (Supplementary Fig. 3), splice sites between sensitivity groups were swapped (Fig. 3), a stronger 3′ss arrangement was employed (Fig. 5) or the strength of the 5′ss was increased (Fig. 6). On the other hand, key features specifically associated with SRRM4 regulation, such as the presence and optimal location of UGC motifs, did not show differential behavior for HS and LS microexons (Fig. 4). Thus, our results are consistent with the model in which the sensitivity to SRRM3/4 is, at least to a large extent, related to the background core splicing configuration of each microexon type (Fig. 1a, Model II), in which (i) HS microexons are much closer than LS microexons to forming productive splicing assemblies in the absence of SRRM4, and (ii) SRRM4’s function on both types of exons is similar.

Next, we developed a mathematical model to further investigate whether the differential sensitivity to SRRM4 could be explained mainly by differences in early spliceosomal recognition, independently of variations in the regulator recruitment or activity. To this end, we developed a minimal mathematical model of splicing and its regulation by SRRM4. Our model (Fig. 7a; see Methods) considers a pre-mRNA that is transcribed and can be bound by the spliceosome (S, considered at a coarse level as a single enzymatic entity, with basal assembly and disassembly rates given by parameters *c_1_* and *c_2_*) as well as the regulator SRRM4 (R, with binding and unbinding rates *c_3_x* and *c_4_*, respectively, where *x* is the regulator concentration). From each bound state, we assume that the exon can be skipped with rate *k_e_* or included with rate *k_i_*. In order to account for the inclusion-enhancing effect of SRRM4, we assume that it enhances spliceosome recruitment by a factor *c_a_*. We consider the model at steady-state, so that the PSI measure corresponds to the ratio of the steady-state concentration of the mature mRNA with the microexon included (I*) over the sum of both the steady state concentrations of included and excluded mature mRNA species (I*+E*). As data to fit the model to, we selected the inclusion profiles of WT LS and HS events, both at the endogenous level and in the MaPSy context, as well as mutations in the 3′ss (BP strong or weak) or in the 5′ss (U1cons). In total, we had 10 groups of sequences, corresponding to the sequences belonging to a given category and sensitivity level, and the corresponding median PSI values at each of the four experimental conditions (GFP and LOW, MID, HIGH expression of SRRM4). We set both the spliceosome and regulator binding rates to 1, in order to keep the number of adjustable model parameters to a minimum, and fitted the remaining parameters (*c*_2_,c_4_*,k_e_,k_i_,c_a_*) as well as the MID and HIGH SRRM4 concentrations relative to the LOW one. To this end, we assessed how well the model could reproduce the median of the inclusion profiles for each experimental condition and sequence group following the assumptions of either Model I or Model II. For Model I, we assume that sequence groups differ only in the parameter corresponding to SRRM4 binding (*c_4_*), whereas for Model II, we assume that sequence groups differ only in the parameter corresponding to spliceosomal recruitment (*c_2_*). The rest of the parameters are assumed to be the same across sequence groups, so that all the dataset must be well fitted by a common set of parameters plus those that are assumed group-specific (either *c_2_* or *c_4_*). We searched for parameter sets using a genetic algorithm, starting from 500 random initial conditions to enhance the chances of finding best-fitting parameter sets. While both models could produce quite good fits to the data, we consistently found the best fits under Model II (Fig. 7b-e and Supplementary Fig. 7a). The reason lies mainly in the GFP condition: some groups of sequences (WT-HS (Fig. 7f), U1cons-HS, BP-stg1-HS (Fig. 7e)) exhibit inclusion in the absence of the regulator, whereas the other groups do not. Model I cannot explain this difference in the absence of the regulator (Supplementary Fig. 7a), whereas this difference can be naturally explained by Model II, which considers that sequence groups may differ in the basal spliceosomal recruitment. Furthermore, we validated that the model cannot fit random data where the values of the different groups of sequences have been permuted within experimental conditions (Supplementary Fig. 7b). Further support to Model II in favor of Model I was provided by the analysis of publicly available SRRM4 PAR-iCLIP data^9^. This data shows that, while the distribution of SRRM4 binding to HS microexons at the 3′ss is narrower (Supplementary Fig. 7c), HS and LS microexons do not differ in their capacity to recruit SRRM4 (Supplementary Fig. 7d,e). These results thus offer orthogonal support to the model where differences in core spliceosomal architecture could be sufficient to produce microexons with different sensitivities to their master regulator.

**Fig. 7:**
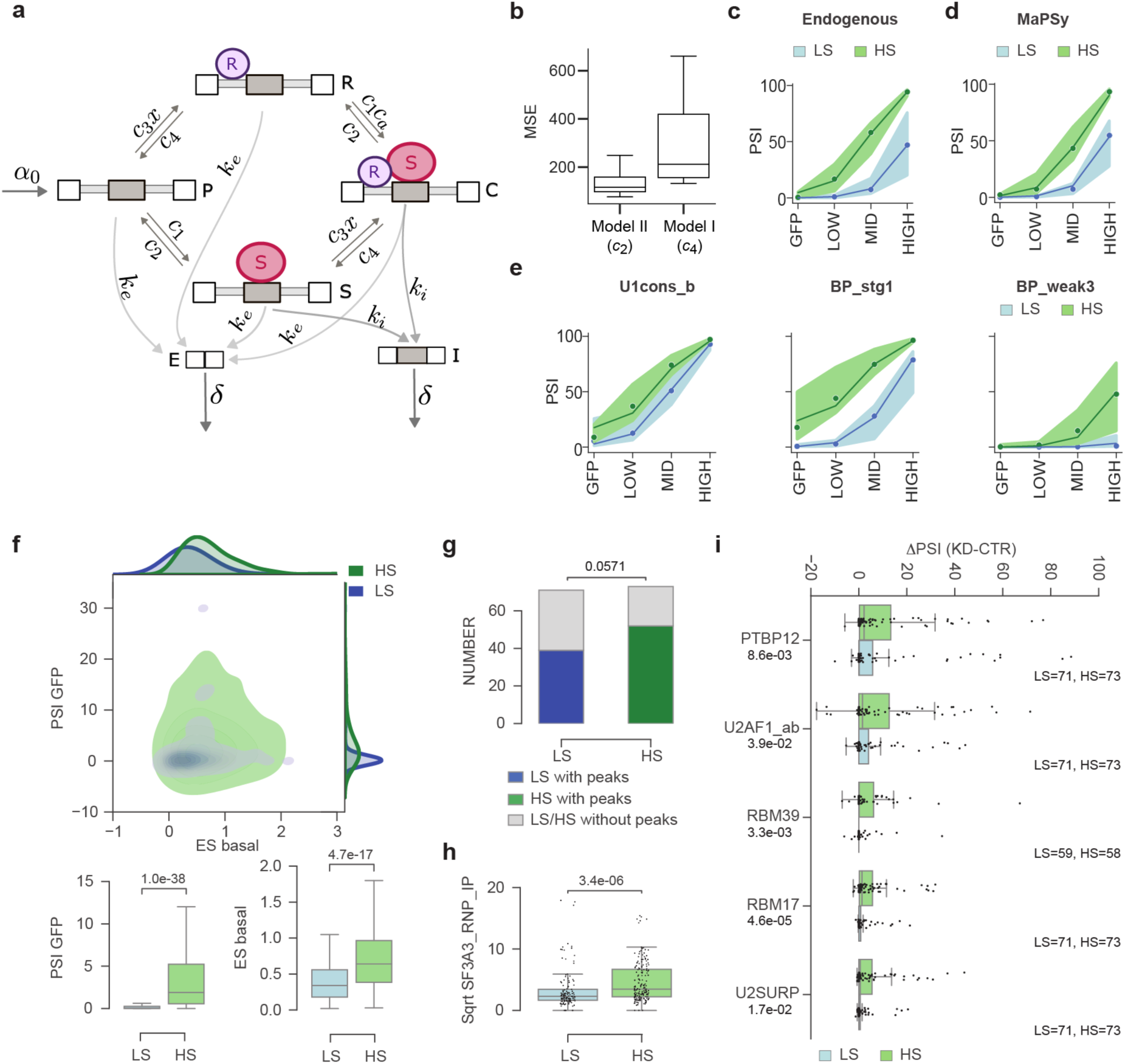
Early recognition of splice sites in the absence of SRRM4 correlates with microexon regulation. (**a**) Integration of the splicing regulation into a mathematical model. (**b**) Distribution of MSE (Mean Squared Error) values for the best parameter sets found in each of 500 independent optimization runs, when assuming groups differ in *c_2_* (Model II) or *c_4_* (Model I). (**c-d**) Mathematical model applied to WT LS (in blue) and HS (in green) events at the endogenous level (c) or in the context of MaPSy (d) in four experimental conditions (GFP or LOW, MID, HIGH level of SRRM4 expression). (**e**) Model applied to mutations in 5′ss (U1cons_b, Fig. 6c) or the 3′ss (BP_stg1, BP_weak3, Fig. 5b). In c-e, the points for each condition represent the medians of the experimental data, the filled area the interquartile range, and the line depicts the model fit. (**f**) Distribution of PSI in control condition (GFP) (y-axis) versus enrichment score (ES) for *in vitro* spliceosomal A complex formation in basal condition (x-axis) for WT sequences for LS and HS events from human and 23 species. The corresponding PSI and ES distributions are separately represented as boxplots below. (**g**) Number of events with or without U2 snRNP related peaks in the 93 nts upstream of the microexons in^57^. Statistics: two-sided Fisher’s Exact test. (**h**) *Ex vivo* U2 snRNP association, represented as square root of SF3A3_RNP_IP peak intensities in^57^, to LS and HS events identified in (f). (**i**) Effects of knocking down individual RNA binding proteins on the splicing pattern of LS and HS events quantified as ΔPSI with respect to control conditions (ΔPSI = KD - CTR). Number of LS and HS events quantified in each condition is reported and each dot represents an event. Statistics in (e,g,h): Mann-Whitney-Wilcoxon test. See also Supplementary Fig. 7.

Additional experimental support to this hypothesis is provided by assays of A complex formation for HS and LS microexons in basal conditions, i.e., with no SRRM4 expression. Specifically, we *in vitro* transcribed the variants bearing the 3′ flanking intronic, microexon and 5′ flanking intronic sequences together with the adaptor sequences, and incubated these RNAs in HeLa cell nuclear extracts under splicing conditions (Supplementary Fig. 7f; see Methods). After separation by electrophoresis, the variants incorporated within the A complex were isolated from the gel and subjected to RNA-seq. The proportion of variants detected in A complex as compared to the proportion in the pool of RNAs used for the reaction was defined as the enrichment score (ES), which is a proxy of the capacity of each variant to recruit early spliceosomal components (Methods). Focusing on the WT human microexons and their orthologs in 23 species, we observed that HS events had significantly higher ESs in basal condition than LS events (Fig. 7f). Similarly, despite being very low in both groups, the levels of inclusion in the absence of SRRM4 (GFP condition) were also significantly higher for HS than for LS exons (Fig. 7f). This significant difference was also observed for all classified endogenous HS and LS events (Supplementary Fig. 7g), a pattern that could not be explained by differences in the strength of their 3′ and 5′ss (Supplementary Fig. 7h,i) but was associated with a differential contribution of ESR/ISR elements (Supplementary Fig. 7j-m).

Next, we took advantage of publicly available data from a recent study mapping the binding of U2 snRNP to 3′ss in the high molecular weight fraction associated with chromatin in HEK 293 cells^57^. Since these cells do not express SRRM3/4, they can be used as a proxy for basal spliceosomal recognition of microexons. Therefore, we asked whether we could detect differential recruitment of U2 snRNP on endogenous HS and LS microexons. For SF3A3, RBM5 and RBM10 IP, a higher proportion of HS microexons presented U2 snRNP peaks in the last 93 nts of the intron upstream to the microexons as compared to LS events (Fig. 7g), and these were of higher intensity (Fig. 7h, Supplementary Fig. 7n-p). Importantly, this did not correlate with differences in mRNA expression of the host genes (Supplementary Fig. 7q) or differences in intensity for other peaks within the same genes (Supplementary Fig. 7r). These data further support the hypothesis that HS and LS microexons are different in their initial basal state, with HS being more primed to enter into the splicing reaction. In line with this, experimental depletion of various splicing factors involved in 3′ss recognition or repression in human cell lines with no SRRM3/4 expression was sufficient to induce mild but significant upregulation of HS microexons but not of LS ones (Fig. 7i).

## DISCUSSION

Since their discovery a decade ago, neural microexons were considered a paradigmatic example of switch-like splicing regulation: expression of their master regulators SRRM3/4, endogenously or ectopically, was sufficient to activate microexon inclusion in any tested cell type^4,6,9,10,12,13,16,58–62^. However, recent work has shown that this switch-like model is incomplete. Two types of SRRM3/4-dependent microexons were identified, one type included in transcripts in endocrine pancreatic cells and neurons, and another expressed only in neurons^4^. Remarkably, this tissue-specific pattern was not driven by co-factors but mainly by the differential sensitivity of each type of microexon to SRRM3/4 activity. While pancreatic microexons were highly sensitive (HS) and needed only low levels of the regulator, neural-specific microexons were lowly sensitive (LS) and required much higher levels of SRRM3/4^4^. Therefore, microexon activation follows a dim-switch like model rather than an on/off one^7^. Interestingly, this opened up the possibility that differences in microexon sensitivity corresponded to different cell type-specific needs. However, it left a major unanswered question: how is microexon sensitivity to SRRM3/4 encoded?

In this study, we aimed at answering this question and understanding the *cis*-regulatory exonic and intronic determinants governing the pattern of microexon splicing and their response to SRRM3/4. We developed a microexon-focused MaPSy approach that recapitulated their regulation by using 93 nts of the 3′ flanking intron - the microexon - 25 nts of the 5′ flanking intron in a minigene reporter and HEK 293 Flp-In T-REx cells expressing either GFP or various SRRM4 levels in an inducible manner. This approach demonstrated the complex contribution of elements in the 3′ss, the 5′ss and the microexons themselves, as well as the conservation of their regulation over ∼450 MYA of vertebrate evolution. Moreover, it reemphasized that exonic length is a major determinant of the regulatory logic of SRRM3/4-dependent microexons. By elongating or shortening individual microexons, we have found that, as a general rule, longer microexons are more efficiently recognized, often to a point where SRRM4 was no longer necessary for their inclusion. This aligns with pioneering work on nSRC^45^ and suggests that the short length of microexons imposes idiosyncratic constraints, ensuring a fine balance between repression in the absence of SRRM3/4 and avoiding excessive repression that would hinder the regulator’s function. We propose that, by adjusting these constraints to the core splicing architecture of each microexon, the optimal dose response level, or sensitivity, to SRRM3/4 is achieved.

To thoroughly assess this possibility, we explicitly confronted two contrasting (yet not mutually exclusive) models that could explain the difference in sensitivity between HS and LS microexons (Fig. 1a). According to the first model (Model I), HS and LS microexons would have a similar “core splicing” architecture (a long BP-AG distance, strong 5′ss, etc.^5,9,11^), but they would differ in their specific SRRM3/4-responding architecture, e.g., by varying in number of UGC motifs, with stronger binding affinities and/or more accessible in the pre-mRNA. This would translate into a stronger response at a given level of SRRM3/4 expression for HS than for LS microexons. In the second model (Model II), the absolute effect of SRRM3/4 regulation would be similar for both HS and LS microexons (i.e., they would be in fact equally sensitive to it), but the “core splicing” architecture of each type would be distinct, with HS microexons being basally more primed for exon inclusion. In both models, the effect of SRRM3/4 at a given concentration can be represented as a enhancing effect towards higher inclusion in a response curve (Fig. 1a). In the first model, such a “push” would be of different magnitude for HS than for LS microexons. In the second model, however, the magnitude of the push would be similar for both types and the difference in resulting PSI would be due to the distinct “starting position” along the inclusion response curve, which is in turn related to the general or core splicing architecture of the microexon. It is crucial to note that for both HS and LS microexons, the default configuration is not sufficient for proper exon recognition and substantial inclusion without SRRM3/4, which is needed to further recruit early spliceosomal components^10^.

Importantly, each model makes specific predictions that we could test. First, in contrast to Model I, Model II predicts that recruitment of SRRM4 should not differ much between HS and LS exons, as we found. Moreover, we find that for both types of microexons, UGC architecture (i.e., the spatial arrangement of the UGC motif relative to other splice site features), when present, seem to be optimized. A second key prediction of Model II is that small changes in core splicing elements (e.g., the addition of ESE/ESSs, strengthening/weakening of the 5′ or 3′ ss, etc.) would be sufficient to cause increases or decreases in microexon sensitivity to SRRM4, both for HS and LS, as observed across all analyses. Third, when such changes in core splicing architecture increase spliceosomal recruitment, i.e., result in a forward shift in the inclusion response curve, they should cause SRRM3/4-independent inclusion for HS but not LS microexons, as we indeed observed. Fourth, similarly, changes in expression in splicing factors involved in splice site recognition or repression in non-SRRM3/4 expressing cells would more often lead to preferential inclusion of HS microexons compared to LS ones. Fifth, early steps of spliceosome assembly should be more readily observed in HS than in LS microexons in the absence of SRRM3/4, as we observed for *in vitro* A complex formation as well as for U2 snRNP recruitment on nascent RNA isolated from chromatin. Moreover, somehow unexpectedly, we also reported the conversion of several CS events into SRRM4-dependent microexons upon exon shortening, whose behavior was usually closer to that of HS microexon. This suggested that the core architecture required for a high sensitivity response to SRRM4 is, to some extent, closer to that of CS exons. Finally, the integration of these concepts into a mathematical model further supported that a different initial state between HS and LS microexons, attributed solely to their basal association with the spliceosome, could be sufficient to explain the patterns of sensitivity we observed in endogenous microexons as well as in our minigene libraries. Therefore, and although we have not formally ruled out Model I and, in fact, it is very possible that it also contributes to microexon sensitivity to SRRM3/4, our data collectively provides strong support for Model II in determining microexon sensitivity. This model is also consistent with the model of global scaling of splicing regulation^25^, by which seemingly small differences in the initial inclusion levels may highly affect the magnitude of the response of exons to splicing perturbations.

Subsequent work should examine the effects of different types of mutations and on a more extended set of microexons, as well as integrate the effects of modulating the activity of other *trans*-acting factors. Moreover, with the exponential growth of machine learning and Artificial Intelligence approaches, complementary analyses of these data should bring us closer to define a microexon code that accurately predicts splicing outcomes under multiple conditions. Finally, whether or not this model for differential sensitivity also applies to the exon targets of other tissue-specific splicing regulators (e.g., RBFOX, NOVA, etc.) needs to be investigated.

### Limitations of the study

First, the experiments presented here were performed in HEK 293 cells. Even though these cells recapitulate the dose response inclusion of the microexons to different levels of SRRM4, they may not fully reconstitute the tissue specific context from neural or pancreatic endocrine cells that could contribute to their finely tuned splicing regulation. Second, the analyses were conducted using a limited subset of microexons from the nested programs (15 LS vs 15 HS), which may not fully capture the diversity of *cis*-acting elements arrangements, particularly in LS microexons with large BP-UGC-AG distances. Likewise, this subset—selected based on their comparable behavior in the MaPSy and endogenous contexts—showed distinct distributions of 5′ss strengths for HS and LS exons, unlike the full set of HS and LS events. Additionally, the pre-mRNA sequences were restricted to 93 nts in the upstream intron and 25 nts in the downstream intron surrounding the microexons and inserted in an ectopic context, which may have led to the loss of longer distance molecular determinants and co-transcriptional splicing regulation. Finally, while the proposed mathematical model shed light into the mechanistic differences between HS and LS microexons and particular relevance of early spliceosomal recruitment, additional experiments are needed to investigate the microexon specific contribution of SRRM4 in assembly and splicing catalysis. As some microexons do not strictly depend on UGC for regulation and that the interplay between the UGC architecture and regulatory activity is complex, it is conceivable that SRRM4 recruitment may occur through alternative mechanisms, such as PPI with spliceosomal components or co-factors. Additionally, we can not rule out that dynamic SRRM4 binding, not captured by current technologies, impacts microexons’ splicing regulation.

## ACKNOWLEDGEMENTS

The authors would like to thank Juan Valcárcel, all members of the Irimia Lab, Lawrence Chasin for providing a comprehensive list of ESRseq scores; Andrey Damianov, Chia-Ho Lin and Douglas Black for providing the HMW peak intensities and discussion on their data (Damianov et al. 2024), Antonio Torres-Mendez for insightful discussion on microexons regulation, Luis Iñiguez-Rabago and Miquel Anglada Girotto for assistance with bioinformatics, David Gray for his insights into pharmacological models, the Genomics Unit at the CRG for assistance with the sequencing, the Protein Technologies Unit at the CRG and the CRG Core Technologies Programme for their support and assistance in this work. Research for this publication has been partially carried out in the Barcelona Collaboratorium.

## AUTHORS CONTRIBUTION

Conceptualization, SoB, SB, RMC, MI; Methodology, SoB, SB, RMC, MI; Software, SB, RMC; Validation, SoB; Formal Analysis, SoB, SB; Investigation, SoB, SB, RMC, MI; Resources, SoB, MI; Writing - Original Draft, SoB, MI; Writing - Review & Editing, SoB, MI; Visualization, SoB, SB, RMC; Supervision, SoB, MI; Project Administration, MI; Funding Acquisition, MI.

## DATA AVAILABILITY

The datasets supporting the conclusions of this article have been deposited to the Gene Expression Omnibus (GSE276143) repository. Source data are provided with this paper.

## CODE AVAILABILITY

All original codes used to generate the results in this manuscript are deposited at the GitHub repositories: https://github.com/simon-bt/lme-mapsy and https://github.com/theobiolab/SRRM4_MapSy_2024.git

The code deposited is meant to allow reproducing the results of the manuscript. However, given its limited annotation and its specificity to the analyses of this paper, it is not meant to be usable for the community for other MaPSy applications.

## SUPPLEMENTAL INFORMATION

Document S1. Supplementary Fig. 1-7 and legends, Minigene reporter sequence. Supplementary Tables 1-21: All tables are provided as standalone excel documents. Source data: Uncropped gels for Fig. 2d,2e,4b,6e,6f and Supplementary Fig. 2j,4c,6f.

## METHODS

### Definition of microexon sensitivities

We have selected microexons previously defined as enriched in pancreas, neural and/or retina samples vs others based on previous studies^4,5,62^. Therefore, these exons tend to be very lowly or not included in most other tissues. We then used RNA-seq data corresponding to (i) the ectopic expression in HEK Flp-In T-REx cells of either GFP (PRJNA474911^10^) or human SRRM4 upon titration after induction using 0 (uninduced, CTR), 0.3 (LOW), 1.2 (MID), 4.8 (HIGH) ng/ml doxycycline (generated in this paper, see below, Supplementary Table 9), and (ii) and the ectopic expression in HeLa FLIp IN cells of either GFP or mouse SRRM3 (PRJNA817274^4^). Microexon inclusion levels (PSI) were quantified for each condition with *vast-tools* v2.5.1^63^. Deprecated events in the final output of *vast-tools* were retrieved from the intermediate raw inclusion tables (option *--keep_raw_incl*). We then defined each event as follows based on various PSI thresholds, irrespectively of the read coverage (Supplementary Fig. 1a, Supplementary Tables 10, 11):

- Non-Responding events: (PSI_HIGH_SRRM4_) – (PSI_GFP_) < 10 AND (PSI_HIGH_SRRM3_) – (PSI_GFP_) < 10.
- Highly Sensitive (HS) events: (PSI_LOW_SRRM4_) – (PSI_GFP_) ≥ 25 OR (PSI_LOW_SRRM3_) – (PSI_GFP_) ≥ 25.
- Lowly Sensitive (LS) events: [(PS_LOW_SRRM4_) – (PSI_GFP_) < 25 OR (PSI_MID_SRRM3_) – (PSI_GFP_) < 25] AND [(PSI_CTR_SRRM4_) – (PSI_GFP_) < 5 OR (PSI_CTR_SRRM3_) – (PSI_GFP_) < 5].

Thirty cryptic (CR) microexons (PSI < 10 in > 90% of the samples with coverage and a PSI range < 25) were manually selected from VastDB to represent different cases with and without UGC and UCUC motifs. Finally, the output was manually curated to assign the sensitivity when NAs were present in any condition.

### mRNA sequencing

RNA was isolated using the RNeasy Mini Qiagen kit and the RNA quality was checked using Bioanalyzer (Agilent). Libraries were prepared at the CRG Genomics Unit using the TruSeq stranded mRNA Library Prep (ref. 20020595, Illumina) according to the manufacturer’s protocol. Each library was sequenced on a HiSeq2500 Illumina machine, to produce an average of 79.6 million 125-nt long paired end reads. This data was submitted to GEO (GSE276143).

### Retrieval and curation of vertebrate microexon orthologs

To generate the evolutionary variants, we first identified the orthologous microexon events in 47 vertebrate species (Supplementary Table 3). For this purpose, we searched LASTZ alignments (available from Ensembl REST API) for available amniote species using as queries each tested human HS and LS microexon and their 93 nts upstream and 25 nts downstream coordinates, −500 and +500 nts respectively. Alignments were reverse complemented if needed and automatically searched for the presence of microexons with maximum 3 nts substitutions and no more than of 40% deletion with respect to the human orthologous event, and AG and GT at 3′ and 5′ ends of the identified exons. Orthologous sequences were then manually curated and checked for completion in the original genome assemblies. For frogs, teleosts and the elephant shark, as well as for additional ammonite species with available “UCSC Chain Files” (https://hgdownload.cse.ucsc.edu/goldenpath/*/liftOver/), we manually searched for each microexon through UCSC liftover and Ensembl Blast searches. The information related to the divergence time per species (million years ago; MYA) and tree were recovered from http://timetree.org/^64^ (Supplementary Table 3). The tree was adapted using *ete3* at http://etetoolkit.org/^65^. Sequences’ alignment provided in Supplementary Fig. 6i was generated using Jalview v2.11.3.3.

### Microexon inclusion and *Srrm3/4* gene expression levels in other species

Orthologous events in *Mus musculus (mm10)*, *Rattus norvegicus (rn6)*, *Gallus gallus (galGal4)*, *Bos taurus* (*bosTau6*) and *Danio rerio* (*danRer10*) were obtained from the “EVENT CONSERVATION” table from *VastDB* (https://vastdb.crg.eu/wiki/Downloads). PSI and gene expression (cRPKM) in tissues from those five species were collected from the “MAIN PSI TABLE” and “MAIN EXPRESSION TABLE” from *VastDB*. Tissues were classified either as Neural, Pancreatic beta cells (when available), Others or discarded (Supplementary Table 12). The PSIs supported by sufficient read coverage (*vast-tools* quality scores VLOW, LOW, OK, SOK) were plotted for all biological samples. The cRPKMs are represented only for tissues in which the PSI of LS and HS events were quantified. Animal silhouettes in Supplementary Fig. 1d were obtained from https://www.phylopic.org/.

### Generation of the barcoded minigene backbone for library cloning

First, a DNM1 minigene encompassing ‘upstream exon (116 nts) - intron (433 nts) - microexon (12 nts) - intron (647 nts) - downstream exon (171 nts) - first 25 nts of the downstream intron’ was constructed for the *DNM1* human microexon (HsaEX0020439). The corresponding genomic region was amplified by PCR from genomic DNA extracted from HEK 293 cells using genElute mammalian genomic DNA miniprep kit (Sigma) and cloned using EcoRV and NotI restriction sites under a *Cytomegalovirus* promoter. Furthermore, PT1 and PT2^66^ sequences were added at the 5′ and 3′ ends of the minigene, respectively, for detection by reverse transcription PCR after transfection (Fig. 1c). Second, for accurate PSI quantification, both the inclusion and exclusion isoforms should be readily identifiable in the output RNA-seq. Therefore, barcodes were added between PT1 and the first exon in the DNM1 minigene backbone prior to the cloning of the variants. For this purpose, the plasmid bearing the DNM1 minigene was amplified by PCR using the Taq Precision Plus enzyme (Agilent Technologies) with primers in PT1 and at the 5′ end of the first exon. An oligonucleotide pool containing the 34-nt long barcode NNNNAGCTNNNNTCAGNNNNTAGCNNNCAGTNNN and flanking sequences was ordered to IDT as ‘hand mix’ ssDNA and amplified through 10 cycles PCR using Taq Precision Plus enzyme (Agilent Technologies). The barcodes were cloned using the Gibson Assembly cloning strategy with a mix of enzymes provided by the CRG Protein Technologies Unit. The plasmids were transformed into Stellar cells (Takara Bio). A pool of an estimate of 209,000 barcoded vectors was obtained. This minigene pool was used as backbone for the cloning of all the libraries used in this study.

### Library design and cloning

A total of 234 events were chosen to build the MaPSy LME (Library of Micro(&)Exons) T0. These included 207 microexons of lengths from 6 to 27 nts with pancreas, neural and/or retina enriched regulation^4,62^ (Supplementary Table 1). Thus, these microexons generally have very low PSIs on most other tissues (Supplementary Fig. 1d, 7g). In addition, it included 27 42-nt long constitutive exons. The microexons included 71 lowly sensitive (LS), 73 highly sensitive (HS) and 63 cryptic/control events. HS and LS microexons encompassed different proportions of pancreatic endocrine cell- and neuronal-specific microexons (Fig. 1c, Supplementary Table 1). The events used in LME T1-4 are listed in Supplementary Table 2 and they were selected based on the similarity between their endogenous and MaPSy-derived inclusion profiles. Specifically, we manually selected events that overall faithfully recapitulated the endogenous response by taking into account several factors: (i) similarity of the magnitude of response in the minigene respect to the endogenous (abs(dPSI_HIGH-GFPendo - dPSI_HIGH-GFPlib)); (ii) response of the minigene in the library in the HIGH condition (dPSI_HIGH- GFP); (iii) response of the minigene in the library in the LOW condition (dPSI_HIGH-GFP: should be as high as possible in HS, and as low as possible in LS events); (iv) low basal inclusion in the minigene in the GFP condition (Supplementary Table 13, see below). In addition, we prioritized a few microexons due to their biological interest.

Each of the five libraries were designed to contain the WT sequences and a subset of the full set of variants (up to 6,000 sequences per library). These variants included various types of mutations and modifications (e.g., UGC mutations, strengthening/weakening of splice sites and other splice signals, semi-deep mutagenesis, inclusion of additional motifs, etc.), as well as swappings between events. All variants and the rationale behind their design are described in detail in Supplementary Tables 5, 14, 15. These designed variants were ordered as ssDNA (from 164 to 200 nt in length) pools to Twist Bioscience (Supplementary Table 16). Each sequence contained 93 nts of the upstream 3′ss, microexon/exon, and 25 nts of the downstream 5′ss, surrounded by two 20-nt long artificial sequences (Mega and Moe). The libraries were cloned into the barcoded DNM1 minigene backbone replacing the central exon and surrounding splice sites (93 nts of the upstream intron and 25 nts of the downstream intron) (Fig. 1c). For this purpose, the plasmid was amplified by PCR using the Taq Precision Plus enzyme (Agilent Technologies) with primers designed to anneal intronic sequences and bear Mega / Moe overhang sequences, and further treated with DpnI (New England Biolabs). The ssDNA libraries were amplified with the Taq Precision Plus polymerase (Agilent Technologies) through 10 cycles PCR and cloned using the Gibson Assembly cloning strategy with a mix of enzymes provided by the CRG Protein Technologies Unit. The resulting plasmids were transformed into Stellar cells (Takara Bio). The total number of clones (barcode-variant pairs) was estimated after transformation to be: LME T0 76,000 clones; LME T1 145,700 clones; LME T2 104,084 clones; LME T3 198,400 clones; LME T4 163,680 clones. The plasmids were amplified and purified (Plasmid Maxiprep Kit, Qiagen).

The final pools of plasmids bearing the variants have the following structure: PT1 BC (34-nt long) exon 1 intron 1 with Mega **VARIANT** (*93 nt-EXON - 25 nt*) Moe intron 2 exon 3 25 nts downstream intron PT2, whose sequences are provided in SI.

### Transfection of libraries, SRRM4 induction and RNA extraction and sequencing

Tetracycline inducible HEK 293 Flp-In T-REx cells expressing either GFP or SRRM4 generated in^10^ were used. Each condition was performed in 6 replicates. Each replicate consisted in a transfection of 6 wells from a 6 wells plate. For each well, 400,000 cells were seeded in 1 ml of culture media the day before transfection and 80 ng of plasmids bearing the barcoded variants were transfected using Lipofectamine 2000 (Invitrogen) following the manufacturer’s instructions. At the time of transfection, doxycycline at various concentrations was added to induce the expression of the protein of interest.

- LME T0: Non-induced GFP line (GFP) and SRRM4 expressing lines treated with 4.8 (LOW); 19.2 (MID) and 50 (HIGH) ng/ml doxycycline were used.
- LME T1/T2: Non-induced GFP line (GFP) and SRRM4 expressing lines treated with 0 (LOW); 1.2 (MID) and 50 (HIGH) ng/ml doxycycline were used.
- LME T3/T4: Non-induced GFP line (GFP) and SRRM4 expressing lines treated with 0 (LOW); 0.032 (MID) and 50 (HIGH) ng/ml doxycycline were used.

For all libraries, 24 hours after transfection, the cells from 6 wells of each condition were pooled and the RNA extracted using the Illustra RNAspin Mini RNA Isolation Kit (GE Healthcare). The consistency of regulation from T1/T2 and T3/T4 is represented as a scatter plot of GFP (VAR-WT) and HIGH (VAR-WT) for the 277 duplicated sequences in Supplementary Fig. 1g.

### Amplification of the inputs and outputs of the libraries for RNA deep-sequencing

In order to associate each barcode with a variant in each plasmid, the pool of plasmids for each library (inputs) was amplified using primers bearing sequences from Illumina (Read 1 and Read 2) and complementary to PT1 and down to Moe (Fig. 1c), represented as a green amplicon). Each input was amplified and sequenced from three independent PCRs [7 cycles each using Taq Precision Plus (Agilent Technologies)]. The pattern of alternative splicing (outputs) was assessed by RT (with oligo-d(T) (IDT) / random hexamer (Roche Diagnostics GmbH) with AMV Reverse Transcriptase (Promega) and 10 cycles PCR with Taq Precision Plus (Agilent Technologies) using six primers (three forward and three reverse) bearing sequences from Illumina for sequencing 5′-Read 1 – (N/NB/NBK) – PT1-3′ and 5′-Read 2 – (N/NB/NBK) – part of last exon of DNM1-3′ reporter (Fig. 1c, represented as a red amplicon). All primers used are reported in Supplementary Table 17.

For both inputs and outputs, PCR elongation times of 5 minutes were used to decrease the likelihood of generating truncated products that could serve as “mega-primer” in the next PCR cycles. The products of amplification were purified on AMPure XP beads (Beckman Coulter). Amplicons were checked using Bioanalyzer (Agilent) and the final steps amplification (consisting in the addition of the Illumina adaptors) were performed at CRG Genomics Unit. Both the input and output PCR products were deep sequenced. After quantification by qubit, amplification was performed by PCR using NEBNext Q5 Hot Start HiFi PCR Master Mix (ref. M0543L) and Index 1 (i7) and Index 2 (i5) primers containing Illumina’s P5 and P7 sequences. Final libraries were analyzed using Agilent Bioanalyzer or Fragment analyzer High Sensitivity assay (ref. 5067-4626 or ref. DNF-474) to estimate the quantity and check size distribution, and were then quantified by qPCR using the KAPA Library Quantification Kit KK4835 (Ref. 07960204001, Roche). Library LME T0 was sequenced in an Illumina HiSeq2000 machine using a 125+125 nt paired end read format; Libraries LME T1/T2 were sequenced in a NextSeq500 machine as 100+200 nt paired end reads; Libraries LME T3/T4 were sequenced in a NextSeq2000 machine as 100+200 nt paired end reads. An average of 13.4 and 37.6 million reads were generated for the inputs and outputs, respectively (Supplementary Table 18). This data was submitted to GEO (GSE276143). For MaPSy T0, each of the six replicates was analyzed individually and the overlap of the six independent replicates is presented in Supplementary Fig. 1f. Based on the high correlation observed, the libraries T1, T2, T3, T4 were also performed in six independent replicates but pooled prior to sequencing [i.e., post amplification and purification on AMPure XP beads (Beckman Coulter)].

### Identification of barcode-variant associations from MaPSy inputs

To identify barcode-variant associations we used the *identify_associations.py*, available at https://github.com/simon-bt/lme-mapsy. Each forward read contains the information of the barcode, whose sequence was retrieved using the regular expression *“.*(CTTGCTCAAC){{s<=5}}(.{{{barcode_length}}})(GAATGTCTAC){{s<=5}}.*”* (PT1 34 nts BC exon 1). Each reverse read contains the information of the associated variant, whose identity was retrieved using the regular expression *’.*(GGGATAAGACGGTAGGC){s<=5}(.{93})’+(.{{{length}}})’+(.{25})(TCGTAGCACGTCACGGTTGG){ s<=5}.*’)* for reads of 200 nts (LME T1-4) and *’(.{{{lenupint-2}}}AG)’+ ‘(.{{{length}}})’+’(GT.{23})(TCGTAGCACGTCACGGTTGGAGCTCCAGCCAGGTTTTCAAGC){s<=7}.*’)’* for reads of different lengths (LME T0), where *lenupint = len(read_sequence) - 42 - 25 - length*, and *length* corresponds to the length of the exon of each variant, and *s* to the number of allowed mismatches in each substring. The sequence identity of each variant was evaluated by calculating the Hamming distance between the retrieved sequence and the reference variant sequence.

Since MaPSy libraries were generated using a pre-barcoded minigene reporter, it is conceivable that the same barcode gets associated with one or multiple variants. Also, it is possible that template switching during PCR amplification leads to the detection of reads supporting spurious barcode-variant associations. To ensure an accurate quantification of variant inclusion, each barcode should be associated with a unique variant, or otherwise filtered out. Filtering of barcode-variant associations was performed using *resolve_associations.py*, available at https://github.com/simon-bt/lme-mapsy. In brief, the identified barcode-variant associations were filtered for maximum percent of the mismatched sequence based on the Hamming distance (*max_mismatch_pct = 50,* for multiple matching variants associated with a barcode we considered the closest matching variant), a minimum read support of 5 for the main association (*min_nreads = 5*), presence of the main barcode-variant association in at least 2 input sequencing replicates (*min_rep = 2*), a maximum of 1 read supporting the second most represented variant associated to a given barcode (*nreads_second_highest = 1*), and maximum percentage (10%) of reads supporting the second most supported variant (*pct_second_best* <= 10). The barcodes that did not meet these requirements to be considered uniquely associated to a variant were considered ambiguous and discarded from further analyses. The set of filtered unique barcode-variant associations for each LME library was used to quantify inclusion of each variant from MaPSy output data (median number of BC/VARIANT is reported in Supplementary Table 16).

### Quantification and correction of variant inclusion levels from MaPSy outputs

The analysis of the output was performed using *quantify_inclusion.py* and *process_inclusion_data.py*, available at https://github.com/simon-bt/lme-mapsy. For each valid barcode supported by at least 3 reads (*min_nreads = 3*), exclusion and inclusion reads in the different experimental conditions were identified using the regular expression *’TGAGCGTGTTGGG(.*?)GCCAGCGAGACCG’*, corresponding to the last and first 13 nts of constant exons 1 and 3, and potentially a sequence in between. Exclusions read counts (EXC) corresponded to those for which no internal sequence was detected (exon1-exon3 junction with no sequence in between). Potential inclusion reads were then further investigated to assess if the sequence between the two flanking exons corresponded to the variant uniquely associated with the barcode. The inclusion reads (INC) were directly accepted and counted if the included sequence matched the reference sequence. We further added a read correction to account for potential INC sequences that do not match the expected variant sequence, and could result from various experimental (template switching) and splicing errors. For this, we calculated a proportion of reads derived from such reads to the sum of total potential inclusion reads (i.e., with any sequence in between exons 1-3). This proportion was used to calculate corrected inclusion level values, such that INC_corrected_ = INC * ((100 + proportion) / 100).

Next, we selected barcode-variant associations supported by at least 3 reads after error correction (min_reads = 3), for which proportion of erroneous inclusion reads was smaller than 75% (max_proportion < 75). For every variant represented by at least 2 barcodes (*min_bc* = 2) in at least 2 samples (min_samples = 2), we derived a single PSI per variant (and condition). For this purpose, we first calculated a corrected PSI value for each barcode-variant association using corrected inclusion reads, such that PSI_corrected_ = * 100 / (INC_corrected_ + EXC). Next, we calculated the median PSI per variant (and condition) of all PSI_corrected_ values (*median_psi*), and an absolute difference between that value and each barcode-variant PSI_corrected_ value (*psi_diff* = *median_psi* - PSI_corrected_) to assess the trend in per-barcode values. We accepted barcode-variant association under the following conditions: (i) if (*median_psi* >= 90 OR *median_psi* < 10) AND *psi_diff* < 10, (ii) elsif (*median_psi* >= 70 OR *median_psi* < 30) AND *psi_diff* < 15, (iii) elsif (30 <= *median_psi* < 70) AND *psi_diff* < 20. For each variant, we then aggregated error-corrected inclusion reads (INC_corrected_) and exclusion reads for filtered barcoded data per variant (and condition) and calculated a single PSI value per variant and condition. Manual inspection of multiple WT constructs show that the PSI values after these two correction steps better recapitulated the endogenous microexon inclusion levels.

Finally, to obtain information for as many designed variants as possible, for those variants for which no PSI was calculated due to the applied thresholds (generally a single barcode associated to the variant), we retrieved the inclusion information, when possible, from intermediate files (CORRECTED_COUNTS, see README.md) and similar steps as above were applied. While these PSIs are less robust, they were included in the analyses to increase the proportion of variants covered and they were nonetheless only a minority of cases. Specifically, for each library and condition, 88.3-96.4% of variants were covered by the main quantification strategy and 3.4-8.7% corresponded to these “rescued” PSIs. The percent of variants per library without an assigned PSI ranged from 0.2% (T4 library) to 3.3% (T1 library).

The level of correlation of the PSI between the initial calculated PSI and the retrieved ones in LME_T0 in condition of HIGH level of SRRM4 expression and six biological replicates are presented in Supplementary Fig. 1f. The PSI for all variants is provided in Supplementary Table 19.

### Specific sequence feature analyses

In Fig. 4a, the most relevant UGC was defined as the UGC mutation having the strongest impact on inclusion levels (negative ΔPSI) in the condition of HIGH level of expression of SRRM4. The most optimal UGC in the “UGC-addition walk” was defined as the single UGC addition having the strongest impact on inclusion levels (positive ΔPSI) in HIGH level of expression of SRRM4. The relative distance between the positions of these two UGCs was calculated for each microexon and the counts plotted as a density. The distances from UGC to AG, Py 3′ end and BP were calculated considering the distances defined for the best predicted BP^67^. In Fig. 6g, features for each variant were recovered using either SVM-BP or MAXENT tools (Supplementary Table 9) and the “variation in feature” was calculated as the score of a feature for a given variant with respect to its corresponding value in the WT sequence. All LS or HS events in LME T1-4 for which the delta feature (VAR-WT) for MAXENT5 (Fig. 6g, left), MAXENT3 (Fig. 6g, right) or agez (Fig. 6g, middle) was not zero were considered for the corresponding analysis after removing the length variants (either elongated or shortened). ESRseq scores^44^ were used to generate the line plots in Supplementary Fig. 2b. The score of each successive hexamer in each of the 15 sequences described in Supplementary Fig. 2a is represented as a line from the 5′ end to the 3′ end of each sequence. The same approach was used to calculate the ESRseq scores for LS, HS, CS and CSrem events upon shortening or lengthening with 15 sequences in LME T1, whose mean values are represented in Supplementary Fig. 2c-f.

In Fig. 7h, publicly available datasets were collected and analyzed using *vast-tools* v2.5.1 (Supplementary Table 11). The change in splicing in the different experimental conditions of LS and HS microexons was calculated as ΔPSI (KD-CTR), irrespectively of read coverage to increase the sample size. For Fig. 7f,g and Supplementary Fig. 7f-h, hg19 coordinates of the LS and HS events described in Supplementary Table 1 were retrieved using liftover (https://genome.ucsc.edu/cgi-bin/hgLiftOver). The intensities of the peaks corresponding to the 3′ss occupancy by U2 snRNP in the high molecular weight chromatin fraction in the last 93 nts of the upstream intron upon immunoprecipitation of RBM5, RBM10 and SF3A3 were retrieved from^57^ (supplementary materials kindly provided by the authors) and used to generate Fig. 7f,g and Supplementary Fig. 7f-h. In addition, to ensure that the results were not biased by differences in gene expression, (i) peak intensities over the entire transcripts with microexons for which U2 snRNP binding was detected, and (ii) their gene expression (*vast-tools* option *--expr*) were analyzed (Supplementary Fig. 7i,j). In Supplementary Fig. 7b-d, PAR-iCLIP tag densities and positions were retrieved from^9^ from the *GSE57278_293TnSR100_PariCLIP_pool2.bedgraph* strand agnostic processed file and matched against hg19 microexon coordinates. Events with tags in the last 200 nts of the upstream intron were retrieved to generate the barplot (Supplementary Fig. 7c) and a distance of 150 nts was chosen to generate the Supplementary Fig. 7b,d.

In Supplementary Fig. 7j bottom, for each sequence variant, we considered the intron + exon + intron sequence and used SpliceAI to predict the probability of each base being a donor or an acceptor site. For this, we followed the instructions in the SpliceAI github repository (https://github.com/Illumina/SpliceAI), FAQ 3. Briefly, we padded the sequence with 5000 “N” at each side, and took the mean of the predictions obtained by each of the five models provided. For each sequence, we checked that the positions with maximum donor or acceptor probability were at the 5’ss or 3’ss, respectively, and used the value at that position for the analysis.

In Supplementary Fig. 7k, the list of ESR and ISR elements were collected from^30,46,68,69^. The number of occurrences of ESR within each exon was counted and divided by the exonic length. The number of ISR^69^ in the 93 nts upstream and 25 nts downstream intronic sequences surrounding each exon was counted and divided by the total intronic length.

### Experimental validations with single minigene transfection

DAAM1 minigene amplified from human genomic DNA, surrounded by PT1/PT2^66^, was cloned under a *Cytomegalovirus* promoter and mutated using primers described in Supplementary Table 17. *apbb1* and *asap1* minigenes from *Danio rerio* were previously generated for^10^. Stronger 5′ss mutations were performed using primers described in Supplementary Table 17. Other variants from the MaPSy were cloned using the same strategy as for the library cloning. Inserts were either ordered as a unique ssDNA encompassing Mega, 93 nts of the 3′ss, microexon, 25 nts of the 5′ss, and Moe, or two overlapping sequences (Supplementary Table 17). The inserts were amplified by PCR using either Mega Fwd and Moe Rev primers or Fwd and Rev primers from the overlapping sequences, respectively. The fragment was cloned using the Gibson cloning strategy into the DNM1 minigene backbone. Specific mutations were generated by PCR around the vector using Taq Precision Plus polymerase (Agilent Technologies). The amplified vector was treated with DpnI (New England Biolabs), T4 DNA polymerase (New England Biolabs) and ligated using T4 DNA ligase (New England Biolabs) following the manufacturer’s instructions. The clones were transformed in *in-house* prepared XL1-Blue heat shock competent cells, single clones were amplified in LB supplemented with ampicillin, purified using Qiaprep miniprep kit (Qiagen) and sequenced (Eurofins).

To investigate the pattern of alternative splicing, the minigenes were transfected. For this purpose, 400,000 cells (HEK 293 or HEK 293 Flp-In T-REx cells expressing SRRM4 in a doxycycline inducible manner) were seeded in a 6 wells plate the day prior the transfection. 24 hours later, HEK 293 Flp-In T-REx cells were transfected with 10 ng of minigene and the expression of SRRM4 was induced with 50 ng/ml of doxycycline. For transient expression, HEK 293 cells were transfected with 10 ng minigene and 1 ug of the plasmid allowing the expression of the protein of interest (beta galactosidase as negative control and SRRM4 human / SRRM234 Amphioxus^10^). The transfection was performed using Lipofectamine 2000, following the instructions of the provider. Cells were collected 24 hours post-transfection and the RNA was isolated using the Illustra RNAspin Mini RNA Isolation Kit (GE Healthcare). The pattern of alternative splicing of individual minigenes was detected by RT-PCR assays. Reverse transcription was performed using oligo-d(T) (IDT) and random hexamer (Roche Diagnostics GmbH) using an enzyme provided by the CRG Protein Technologies Unit in 5X First-Strand Buffer (250 mM Tris-HCl pH 8.3; 375 mM KCl; 15 mM MgCl_2_). The PCR was performed using primers PT1 and in the last exon of the minigene with the enzyme GoTaq (Promega). The amplicons were separated on 7% acrylamide 37,5:1 (PanReac Applichem) run in TBE 1X (10.8 g Trizma base; 5.5 g Boric Acid and 4 ml of 0.5M EDTA pH 8.0 per liter) and detected by Gel Red staining (Biotum) using the gel doc software Quantity One 4.6.9. Fiji (version 1) was used for the quantification of the amplicons’ intensities. The PSI for each biological replicate is calculated as 100 * [inclusion / (inclusion + skipping)] and the quantifications are provided in Supplementary Table 20.

### *In vitro* spliceosomal A complex formation analyses

The ssDNA libraries were converted into dsRNA by PCR using primers in T7 promoter _ Mega Fwd and Moe Rev sequences and the Taq Precision Plus enzyme (Agilent Technologies). Cy5-CTP/Cy5-UTP (Cytiva) labeled RNA were transcribed from the PCR templates using the Megascript T7 Transcription kit (Ambion) according to the manufacturer’s instructions. Each library was assembled in four independent reactions. Per reaction, 15 ng/ul fluorescently labeled RNA were incubated with 3 microliters of HeLa cell nuclear extracts (CILBIOTECH, CC-01-20-50) supplemented with 3 mM MgCl_2_, 24.9 mM KCl, 3.33% PVA, 13.3 mM HEPES pH 8, 0.13 mM EDTA,13.3 % glycerol, 0.03 % NP-40, 0.66 mM DTT, 2 mM ATP and 22 mM creatine phosphate in a final volume of 9 microliters. The mixture was incubated for 18 minutes at 30°C. 1 microliter of heparin (50 ug/ul stock) was added and incubated for 10 minutes at room temperature. 3 microliters of 50% glycerol were added and 9 microliters loaded on a composite gel (4% acrylamide, 0.05% bis-acrylamide, 0.5% agarose, 50 mM Tris, 50 mM Glycine). The gel was run for ∼ 4 hours at 200 Volts in 50 mM Tris / 50 mM Glycine buffer. After electrophoresis, the fluorescence was detected using a Typhoon PhosphorImager (Amersham and corresponding Cytiva software). The pieces of gel containing the products corresponding to twice two reactions of A complex formation were cut (Supplementary Fig. 7f) and the RNA extracted in an RNA elution buffer (0.5 M NH_4_Ac, 10 mM MgCl_2_, 1 mM EDTA, 0.1 % SDS) for 6 hours under shaking at 50 degrees. The supernatant was extracted with phenol / chloroform and precipitated with three volumes of ethanol absolute. The RNA was resuspended in water and used for one RT reaction further used in 2 PCR. The reverse transcription was performed using the Moe Rev primer using the AMV Reverse Transcriptase (Promega). The PCR was performed using the Taq Precision Plus (Agilent Technologies) and primers bearing Illumina compatible reads, the corresponding amplicons are represented in green (Supplementary Fig. 7b). The PCRs were pooled prior to purification on AMPure XP beads (Beckman Coulter). In parallel, the RNA used to perform the assay were amplified by RT-PCR to determine the enrichment scores of each variant with respect to its initial expression. The enrichment scores (ES) for libraries T1, T2 are provided in Supplementary Table 21. Each library was sequenced on an Illumina NextSeq500 machine, to produce an average of 1.95 and 1.56 million 100+200 long paired end reads for inputs and outputs, respectively. This data was submitted to GEO (GSE276143). Spliceosomal A complex formation on ITSN1 was performed as described in^10^ on substrates bearing 93 nts of the flanking 3′ss _ microexon _ 25 nts of the flanking 5′ss surrounded by Mega and Moe sequences. Quantification of the biological replicates presented in Fig. 6f is provided in Supplementary Table 8.

### Mathematical model of microexon regulation by SRRM4

We modeled microexon inclusion regulation by SRRM4 with a system of coupled Ordinary Differential Equations (ODEs). See Fig. 7a for a cartoon of the model with the species and rates. The model considers an unbound pre-mRNA species *P* which is transcribed at rate α_0_. The spliceosome, considered at a coarse level as a single enzyme, assembles around the microexon on the pre-mRNA at rate *c_1_* to yield species S, and disassembles at rate *c_2_*. On the other hand, SRRM4 can bind to the pre-mRNA at rate *c_3_x*, with *x* the concentration of SRRM4, to yield species *R*, and unbinding occurs at rate *c_4_*. We assume that bound SRRM4 can favor spliceosomal recruitment to the microexon, by enhancing its assembly by a factor *c_a_*>1. We also assume that SRRM4 binding can follow spliceosomal assembly, and for simplicity we assume that this happens at the same rates as when there is no spliceosome bound.

In addition to binding and unbinding, the pre-mRNA can undergo microexon skipping or inclusion. We assume that there is a skipping rate *k_e_*, and the inclusion rate when the spliceosome is present is *k*_i_. Finally, the mature mRNA products are degraded at rate δ. This yields the following system of coupled ODEs:

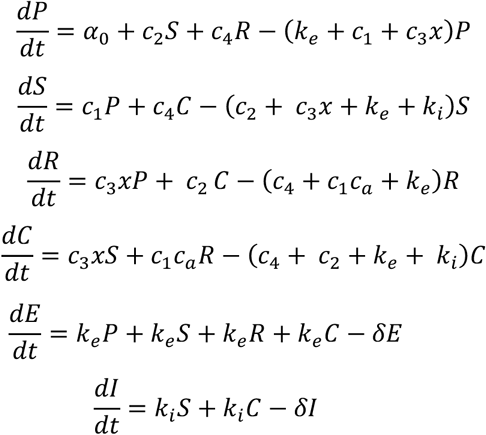

PSI is quantified as: 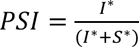, where Y* denotes steady-state species Y.

It can be shown that PSI is independent of *α*_0_ and δ. Therefore, we set both parameter values to 1, without affecting the results. In addition, we set *c_1_*=1 (time unit)^−1^ and *c_3_* = 1 (time unit *x* concentration unit)^−1^, so that spliceosomal and regulator affinity/recruitment are only controlled by a single parameter each (*c_2_* and *c_4_*, respectively). This allows keeping the number of free parameters low while still allowing to parameterise spliceosome and regulator affinity/assembly efficiency. In order to find the steady state, we solve for the solution of the corresponding linear system using the linalg.solve function from the Python numpy package. We double checked that the same solutions are obtained by integrating the ODE system until steady-state using the *odeint* function of Scipy. The corresponding code is provided at https://github.com/theobiolab/SRRM4_MapSy_2024.git.

### Model fitting to experimental data

In order to test whether the model could account for the diversity of the responses to SRRM4 observed in the data, we selected the responses of the WT LS and HS for endogenous microexons and the minigene constructs, as well as the mutants that make the interaction with the U1 snRNA stronger (U1cons_b), and the mutants that make the BP stronger (BP_stg1) or weaker (BP_weak3). We took the median of the PSI values across the sequences for the various groups at each of the four regulator concentrations (GFP, LOW, MID, HIGH). We fixed the concentration of SRRM4 to 0 for the GFP condition, and to 1 concentration unit for the LOW condition, so that the concentrations of the MID and HIGH conditions are in units of SRRM4 concentration in the LOW condition. We then searched for parameter sets that would best fit the data allowing the groups to share the MID and HIGH SRRM4 concentration, the skipping *(k_e_)* and inclusion *(k_i_)* rates, and the regulator effect *c_a_*. When testing Model I, we allowed groups to differ in *c_4_*, with *c_2_* shared, whereas when testing Model II, we allowed groups to differ in *c_2_*, with *c_4_* shared.

Given the coarse level of our models and the lack of measurements for the parameters, we allowed the parameters to be within a broad range, spanning from 10^−6^ to 10^6^, except for *c_a_*, which was restricted to the interval [1,10^6^] to account for an enhancing effect of the regulator on spliceosomal recruitment. We searched for optimum parameter sets by using the genetic algorithm implementation of the *eaSimple* function of the Python Deap package for evolutionary algorithms. We used the following settings and hyperparameters: simulated binary crossover with eta=0.25 and probability of crossover 0.25. Tournament selection with size 3, gaussian mutation, a population size of 400, 300 generations, probability of mutating an individual 0.5, the standard deviation of the gaussian mutation 0.25, and the probability of mutating an individual parameter 0.5. After this, the best solution was refined by refining the parameter allowed to change among groups by local minimization, using the minimize function of *scipy* optimize, starting from the identified parameter value as well as 10 additional random conditions from within the bounds. The genetic algorithm was set to minimize the mean square error between the median of the data and the model predictions per sequence group, defined as, the squared of the differences between the median data PSI and the model PSI for each of the four experimental conditions, averaged across the four conditions, and added over the 10 sequence groups (WT-LS, WT-HS, WT_MaPSy-LS, WT_MaPSy-HS, U1cons-LS, U1cons-HS, BP_stg-LS, BP_stg-HS, BP_weak-LS, BP_weak-HS). In addition, the parameters corresponding to the SRRM4 levels were forced to be increasing between the LOW, MID and HIGH conditions by adding a penalty of 10000 to the error function if the values were decreasing.

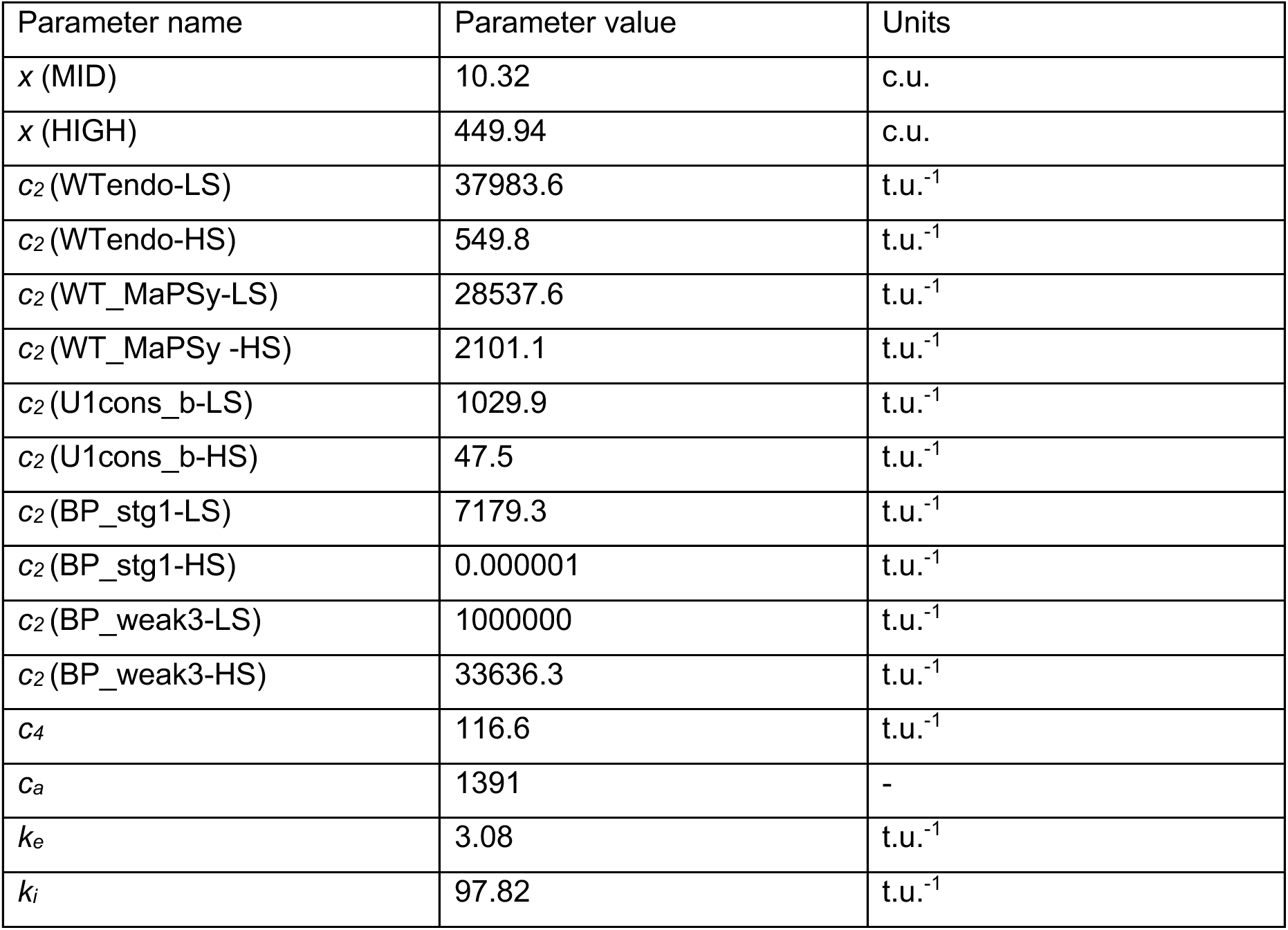

**Parameter values** for the modeling plots in Fig. 7d.

c.u: Concentration of SRRM4 in the LOW condition (*x* (LOW)=1 c.u.)

t.u: (arbitrary) time units.

### Model fitting to random data

To verify that the model cannot fit random data, we generated 9 random datasets, each with 10 trends obtained by randomly permuting the median values of the data within each experimental condition (that is, we permuted within columns if we represent the data as a matrix of 10 rows (groups of sequences) and 4 columns (GFP, LOW, MID, HIGH). For each dataset, we run the same optimization procedure as described above but allowing the parameters to vary even within broader ranges (10^−8^ to 10^8^) to ensure the parameter range was not the limiting factor. We treated each of the 10 random trends as corresponding to the trend of a group of sequences, and therefore having a unique *c_2_* or *c_4_* depending on the model allowed.

### Data visualization

In all panels with boxplots, the boxes represent the interquartile range (IQR) of the PSI or ΔPSI from the 25^th^ percentile to the 75^th^ percentile, the median is represented by the line in the box, the whiskers represent the minimal and maximal values within a 1.5 times the IQR from the 25^th^ and 75^th^ percentile. Any data points beyond this range are not displayed. Line plots were generated using the sns.lineplot() function from the Seaborn library. The central tendency is represented by the mean, and the shaded region corresponds to the 95% confidence interval, estimated through 1000 bootstraps. In Fig. 7c-e and supplementary Fig. 7a, the central tendency is the median of the data, the line corresponds to the model prediction, and the shaded region to the data IQR.

