## Supplementary Figures and Table Legends for "Core splicing architecture and early spliceosomal recognition determine microexon sensitivity to SRRM3/4"

**SUPPLEMENTAL FIGURES**

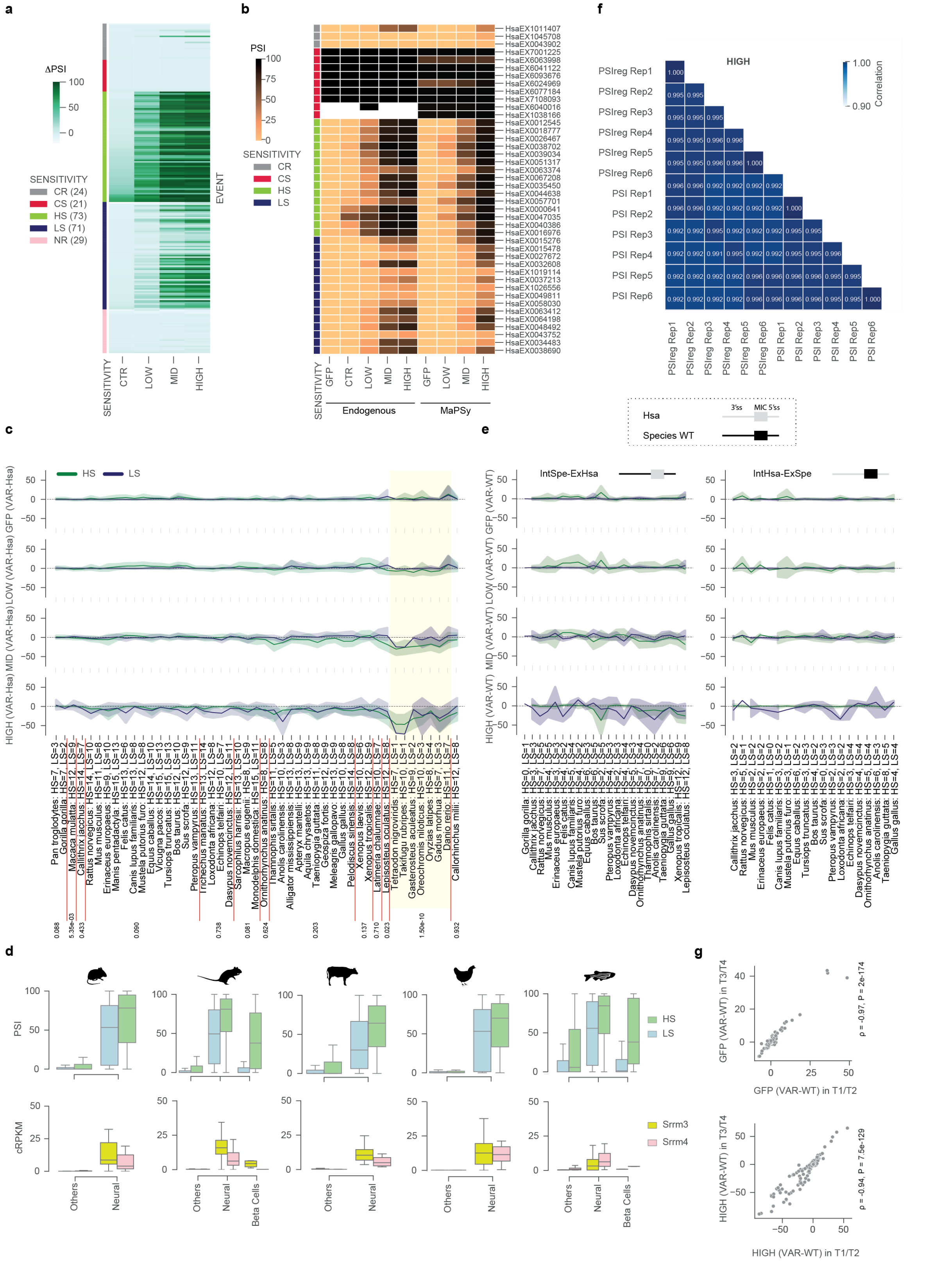

**Supplementary Fig. 1: Microexon sensitivity to SRRM4 expression is largely conserved from shark to human.**

(**a**) Different microexon groups were defined according to their response upon titration of SRRM4. Supervised heatmap representing the ΔPSI for each event (rows) under different conditions (columns) with respect to cells expressing GFP. Conditions: HEK 293 Flp-In T-REx cells expressing SRRM4 in a doxycycline dependent manner, CTR are cells not treated with doxycycline (only leaky expression), while LOW, MID, HIGH are cells treated with increasing concentrations of doxycycline (Methods). The membership of the events to either LS/HS/CR/CS/NR microexons is indicated by the colored squares on the left side of the heatmap. The number of events for each category is shown within brackets. (**b**) Heatmap showing the pattern of splicing of the selected events both in the endogenous and in the MaPSy context (represented as the median of PSI values in the different libraries in which the events were quantified). White squares correspond to missing values due to insufficient read coverage. (**c**) The lines represent an estimate of the central tendency and the corresponding 95% confidence interval of the ΔPSI (VAR-Hsa) of orthologous WT sequences from all tested species with respect to human (Hsa) in GFP and LOW, MID, HIGH expression of SRRM4 conditions for HS (green) and LS (blue) events accordingly to the defined sensitivity in human (Fig. 1). The species are indicated at the bottom as well as the number of HS and LS events represented per species. Teleosts are highlighted using a yellow background as in Fig. 1e**.** The red lines separate the species for each phylogenetic node. The numbers at the bottom show the results of two-sided Mann-Whitney tests between the corresponding node and all other ones combined for the condition HIGH(VAR-WT). (**d**) Top: Distribution of PSIs in various tissues for orthologous of LS (blue) and HS (green) events from *Mus musculus*, *Rattus norvegicus*, *Bos taurus*, *Gallus gallus* and *Danio rerio*. Bottom: mRNA expression levels (cRPKMs) of *Srrm3* (yellow) and *Srrm4* (pink) in each species. (**e**) Schematics of the sequences involved in each swapping experiment. Intronic sequences from either a human event (grey) or another species (black) were swapped to generate the chimeric constructs depicted at the top of each subpanel, where the exonic part is either from human or from another species, respectively. The lines represent an estimate of the central tendency and the corresponding 95% confidence interval of ΔPSI (VAR-WT) between the variants in which the microexon sequence is from human and both flanking introns are from its ortholog in the indicated species (left), or in which the microexon sequence is from a given species and both flanking introns are from its human ortholog (right). Species involved in the shuffling are listed along the x-axis together with the number of HS and LS events represented per species. (**f**) Correlation of PSI between 6 biological replicates in the condition of HIGH expression level of SRRM4. PSI* correspond to the final output of the quantification pipeline while PSIreg* values are recovered from an intermediate file (see Methods). ‘Rep’ indicates each of the 6 replicates. (**g**) Correlation of the 277 sequences present in both T1|T2 and T3|T4 in four experimental conditions (GFP and LOW, MID, HIGH expression of SRRM4).

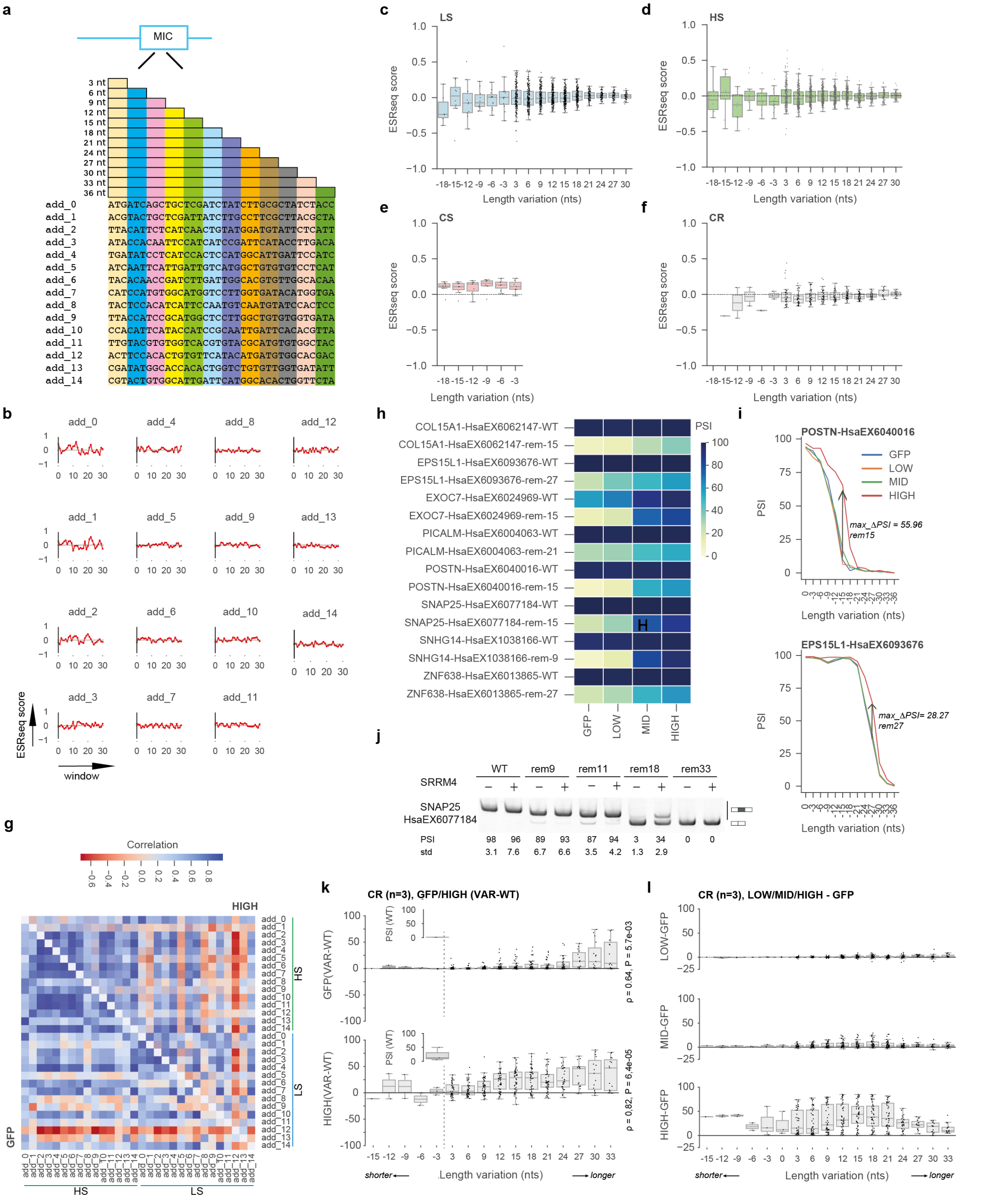


**Supplementary Fig. 2: Exon length and exonic *cis*-acting elements contribute to SRRM4 mediated splicing regulation.**

(**a**) Scheme of the strategy used to increase the length of the microexons. For each microexon, sequences from 3 to 36 nucleotides were designed and added in the middle of the microexon by increments of 3 nucleotides (color coded). The sequences used are provided. (**b**) ESRseq scores provided for each hexamer with a 1-nt sliding window[44] for each of the 15 sequences used in Fig. 2. Scores range from -1 to 1 depending on whether they are likely to harbor negative or positive regulatory motifs, respectively. Scores close to 0 are less likely to have splicing regulatory elements. (**c-f**) ESRseq scores obtained upon length variation of LS, HS, CS and CR events represented in Fig. 2a-c, Supplementary Fig. 2j,k. (**g**) Pearson correlation matrices representing the impact of each designed sequence when added into LS and HS microexons under two experimental conditions (GFP lower triangle, HIGH upper triangle), quantified as ΔPSI (VAR-WT). (**h**) PSI across four experimental conditions (GFP vs LOW, MID, HIGH expression of SRRM4) for each CS WT (42 nts) and their most SRRM4-responding shortened variants (in which the indicated number of nucleotides N have been removed, remN). (**i**) Two examples of CS events whose shortening results in SRRM4-responding variants (EPS15L1-HsaEX6093676 and POSTN-HsaEX6040016). The arrow indicates the maximal value of ΔPSI (HIGH-GFP). (**j**) RT-PCR assays showing the splicing patterns of various SNAP25-HsaEX6077184 splicing reporters (WT or following the removal of either 9, 11, 18 or 33 nucleotides) under control condition (-) or upon expression of SRRM4 (+) in HEK 293 cells. The PCR amplicons corresponding to inclusion / skipping are indicated with the squares. PSI and standard deviations (std) from at least three biological replicates per minigene variant are provided. (**k**) Change in CR exon inclusion upon shortening or lengthening was quantified as ΔPSI (VAR-WT) in either the GFP condition or under HIGH level of expression of SRRM4. Each data point corresponds to the effect of adding a given number of nucleotides from one of the fifteen designed sequences (positive number on the x-axis), or removal of a given number of nucleotides from the exonic sequence (negative number on the x-axis). The length of the WT is indicated by the dashed vertical line. The numbers at the right side of each subpanel indicate the Spearman correlation (ρ) and corresponding p-value (*P*) of the effects in PSI or ΔPSI according to length variation. (**l**) Impact of length variation on the pattern of splicing and regulation under different levels of expression of SRRM4 with respect to GFP condition for each sequence, expressed as ΔPSI (HIGH/MID/LOW-GFP) for CR events.

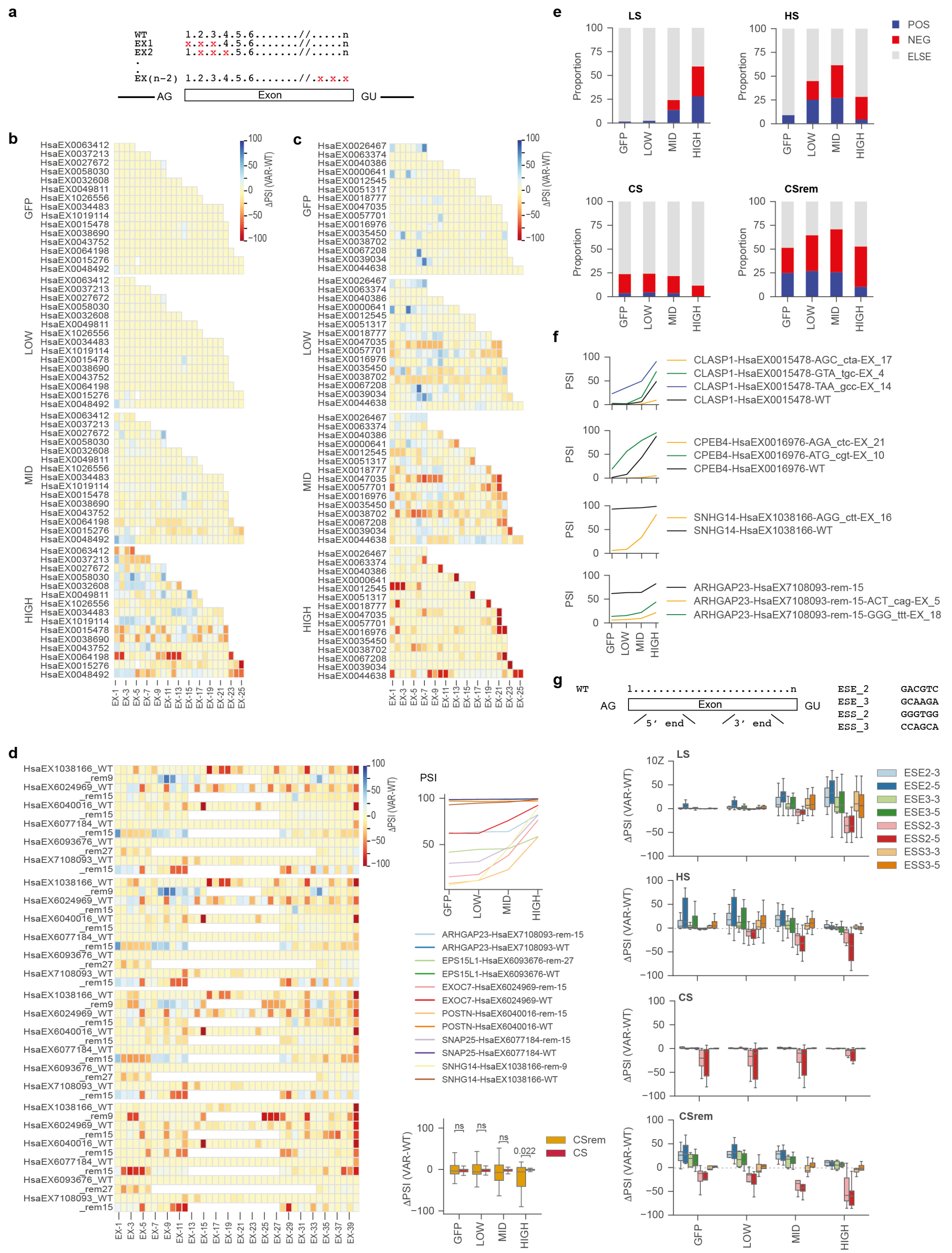


**Supplementary Fig. 3: Exonic elements contribute to splicing and regulation by SRRM4.**

(**a**) Approach used to perform a semi-deep mutagenesis scan of the exon. Three successive nucleotides (indicated in red) were mutated at the time using 1-nt sliding windows and following a systematic pattern of nucleotide substitution: A > C, G > T, C > A, T > G. (**b-d**) For each event and experimental condition (y-axis), the ΔPSI (VAR-WT) upon mutation at different exonic positions (x-axis) is shown as a heatmap (blue/red represents higher/lower inclusion than the WT) (LS in (b) and HS in (c)). For CS events (d), the positions of the mutations in the short exonic version have been aligned to their corresponding WT version of 42 nts (note that because of the removal, the central nucleotides are different). On the right side of panel (d), the line plots represent the PSI in the same experimental conditions for each of the CS events and their corresponding shortened variants. The boxes show the quantification of the perturbations observed on the heatmap. Statistics: Mann-Whitney-Wilcoxon test two-sided with Bonferroni correction (ns: non significant). (**e**) Stacked bar plots for LS, HS, CS and CSrem events in the four experimental conditions counting the number of mutations leading to ΔPSI (VAR-WT) > 10 (POS), ΔPSI (VAR-WT) < -10 (NEG) or without effect (ELSE). (**f**) Examples of changes in PSI and SRRM4 response for variants of a LS (CLASP1-HsaEX0015478), a HS (CPEB4-HsaEX0016976), a CS (SNHG14-HsaEX1038166) and a shortened CS (ARHGAP23-HsaEX7108093-rem-15) event. (**g**) Top: Representation of the approach used to introduce ESR hexamers within the exon. Two ESEs and two ESSs were inserted either at the 5′ end (from position 4 to 9) or at the 3′ end (from position n-9 to n-4, n being the exon length) of the exon as indicated. The sequences of each hexamer are shown (Supplementary Table 7). Bottom: Distributions of ΔPSI (VAR-WT) in GFP vs LOW, MID, HIGH expression of SRRM4 for LS, HS, CS and their corresponding CSrem events. The colors indicate both the hexamer and the position where it was inserted.

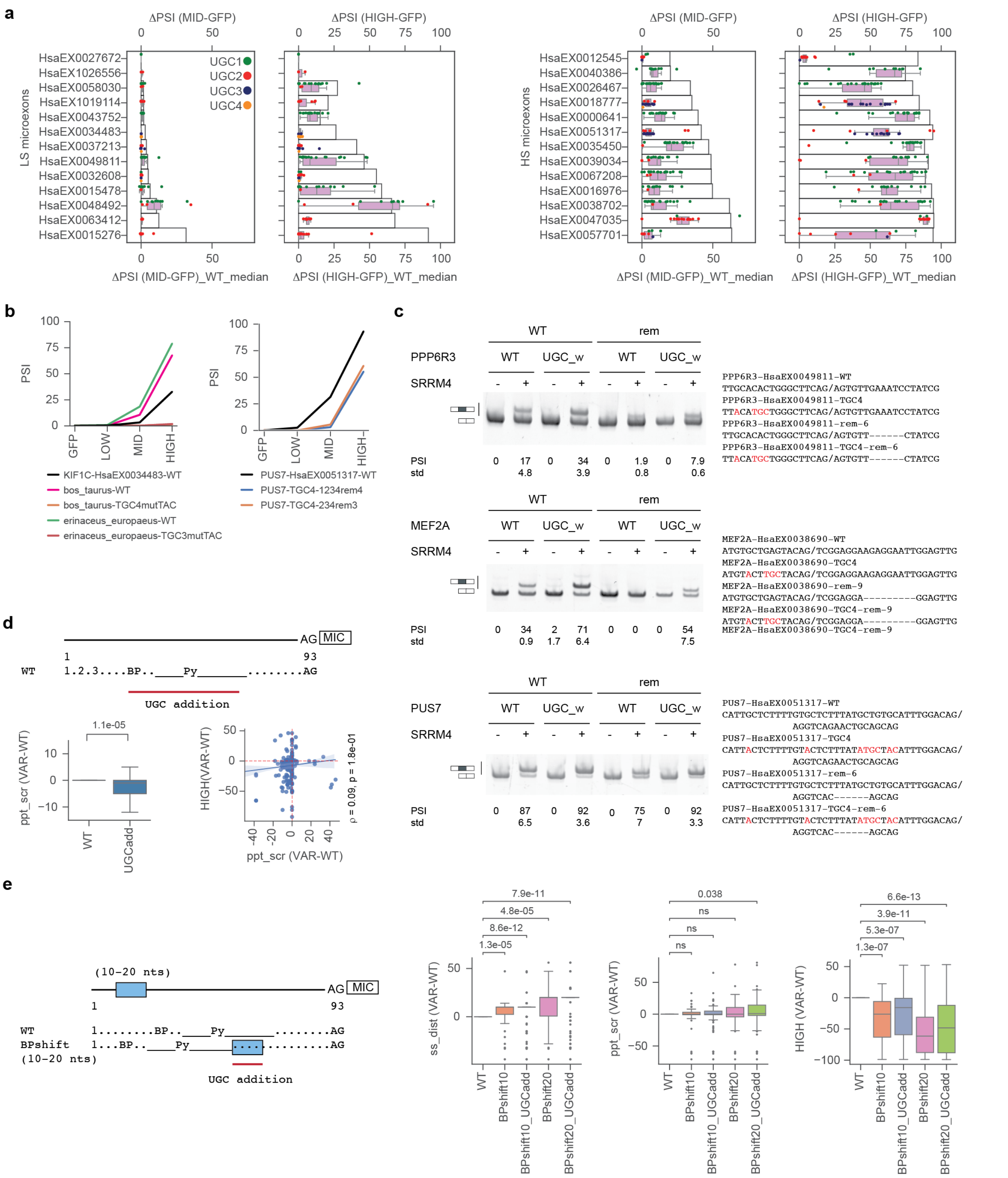


**Supplementary Fig. 4: Effect of UGC mutation and insertion on splicing regulation of LS and HS events.**

(**a**) Effect on the response to SRRM4 at MID and HIGH conditions of mutating all UGCs in the orthologous variants. The barplots represent the median ΔPSI for all the orthologous WT constructs in either MID or HIGH SRRM4 expression conditions with respect to GFP. Boxplots show the distribution of the ΔPSI of the UGC mutant variants, with each dot representing a UGC variant from a given species. The total number of UGCs considered for the mutagenesis for each event are indicated by the color of the dots. (**b**) MaPSy-derived PSI values under four experimental conditions (GFP or LOW, MID, HIGH expression of SRRM4) for selected variants of the KIF1C and PUS7 minigenes tested in Fig. 4b. **(c)** RT-PCR assays showing the splicing pattern of different variants (WT or mutation from the "UGC-addition walk" combined or not with exon length shortening) of three events under control condition (-) or expression of human SRRM4 (Hsa). PSI and standard deviations (std) from at least three biological replicates are provided. (**d**) Top: Schematics of the location at which 1, 2 or 3 UGCs have been added in the context of the WT sequence. Bottom left: Variation of the *ppt_scr* of each variant with respect to its corresponding WT upon addition of 1, 2 or 3 UGCs downstream to the best predicted BP. Outliers were removed for visibility. Statistics: Mann-Whitney-Wilcoxon test. Bottom right: Correlation between the changes in *ppt_scr* and the variation in PSI in the HIGH condition from variant to WT. Statistics: Pearson correlation and p-value are provided. (**e**) Top: Location at which 1, 2 or 3 UGCs were added in the context of a modified WT sequence in which the BP and Py tract were displaced together either 10 or 20 nts upstream to their initial position. Bottom: Variation of *ss_dist*, *ppt_scr* and PSI in the HIGH condition of each variant with respect to its corresponding WT upon addition of 1, 2 or 3 UGCs in the 10 or 20 nts downstream to the BP and its associated Py Statistics: Mann-Whitney-Wilcoxon test two-sided with Bonferroni correction (ns: non significant).

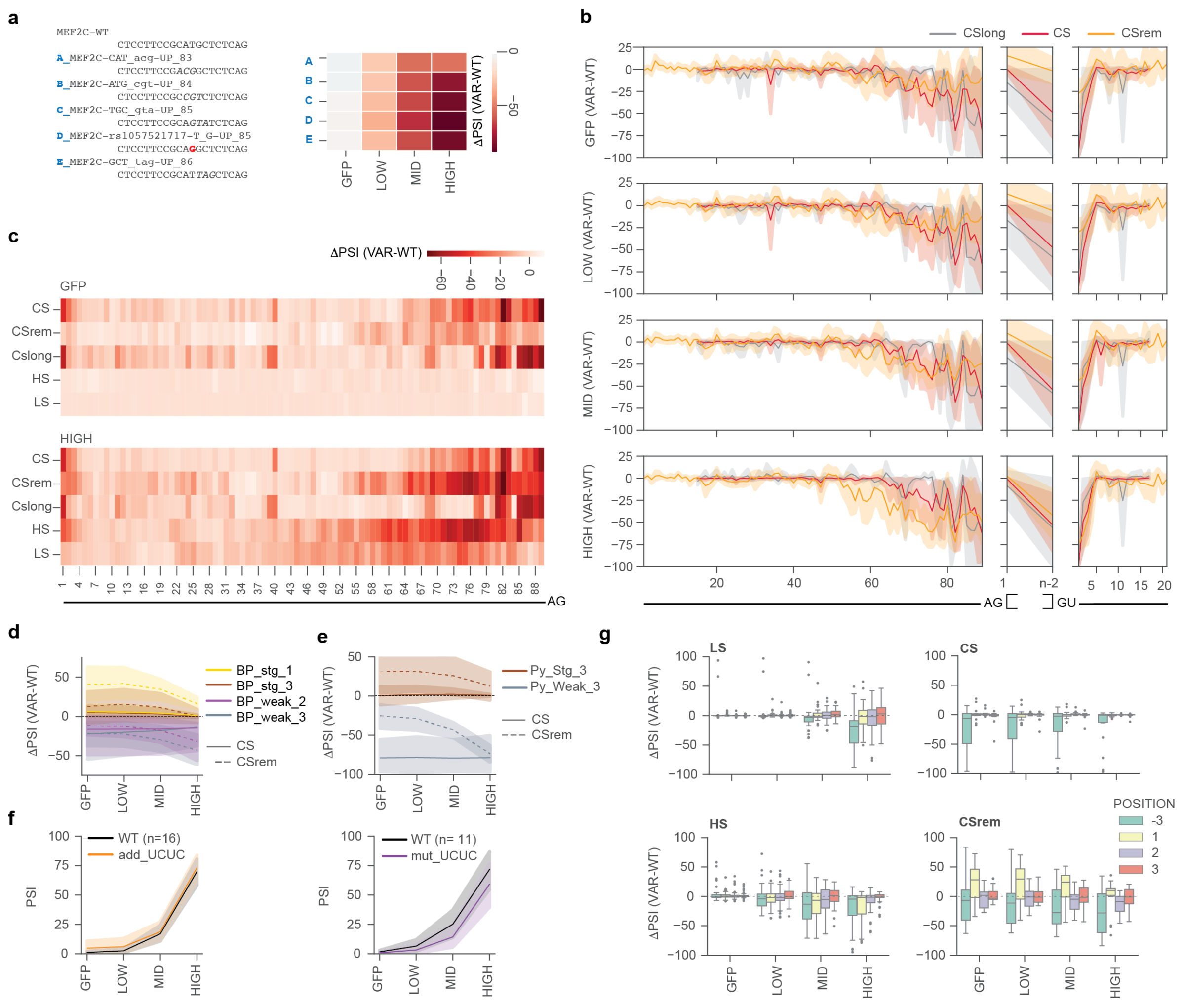

**Supplementary Fig. 5: Intronic core splicing elements are relevant for regulation by SRRM4.** (**a**) The heatmap depicts the effect of mutations with respect to WT [ΔPSI (VAR-WT)] in four experimental conditions (GFP and LOW, MID, HIGH expression of SRRM4) in the 3′ ss of MEF2C-HsaEX0038702 at positions overlapping the SNP (rs1057521717 T>C). Sequences of the last 20 nts of the 3′ss of the WT and the different variants are reported below the heatmap, with the SNP indicated in red. (**b**) Effect of mutagenesis scan as in Fig. 5a for CS (red), their corresponding short version (CSrem, orange) and other CS (CSlong, grey) events. For the exonic sequence, only the impact of mutating the first (1) or last three exonic nucleotides (n-2) is reported (full set in Supplementary Fig. 3d). The positions of the mutations are indicated along the x-axis according to the nomenclature described in Fig. 5a. (**c**) The heatmaps depict the ΔPSI (VAR-WT) upon mutations (Fig. 5a and Supplementary Fig. 5a) in the 3′ss for LS, HS, CS, their shorter versions (CSrem), and other CS (CSlong) events in conditions of expression of GFP or HIGH level of SRRM4. The positions of the mutations are indicated along the x-axis according to the nomenclature described in Fig. 5a. (**d,e**) The lines represent an estimate of the central tendency and the corresponding 95% confidence interval of the ΔPSI (VAR-WT) between BP variants (d) or Py variants (e) and their corresponding WT in four experimental conditions (GFP vs LOW, MID, HIGH expression of SRRM4). In d, the best predicted BP was mutated to either 2 strong (stg) or 2 weak BP motifs. Plain lines represent the impact of the mutations on CS events while the dashed lines represent the effect on their corresponding short versions. (**f**) The lines represent an estimate of the central tendency and the corresponding 95% confidence interval of the PSI in four experimental conditions (GFP vs LOW, MID, HIGH expression of SRRM4) for both LS and HS WT constructs or the corresponding variants in which a UCUC motif was added to the WT sequence when absent (left) or removed when present (right). The total number of events for each type of mutation (n) is indicated in the legend. No significant difference between WT and mutated PSI distributions was observed for any condition (Mann-Whitney-Wilcoxon test). (**g**) Distributions of the ΔPSI (VAR-WT) in GFP vs MID, HIGH expression of SRRM4 between variants and their corresponding WT for either LS (top left), HS (top right), CS (bottom left) and their corresponding short versions (bottom right) events. Variants include deep mutagenesis at position -3 in the 3′ss and positions 1, 2, 3 at the 5′ end of the exon.

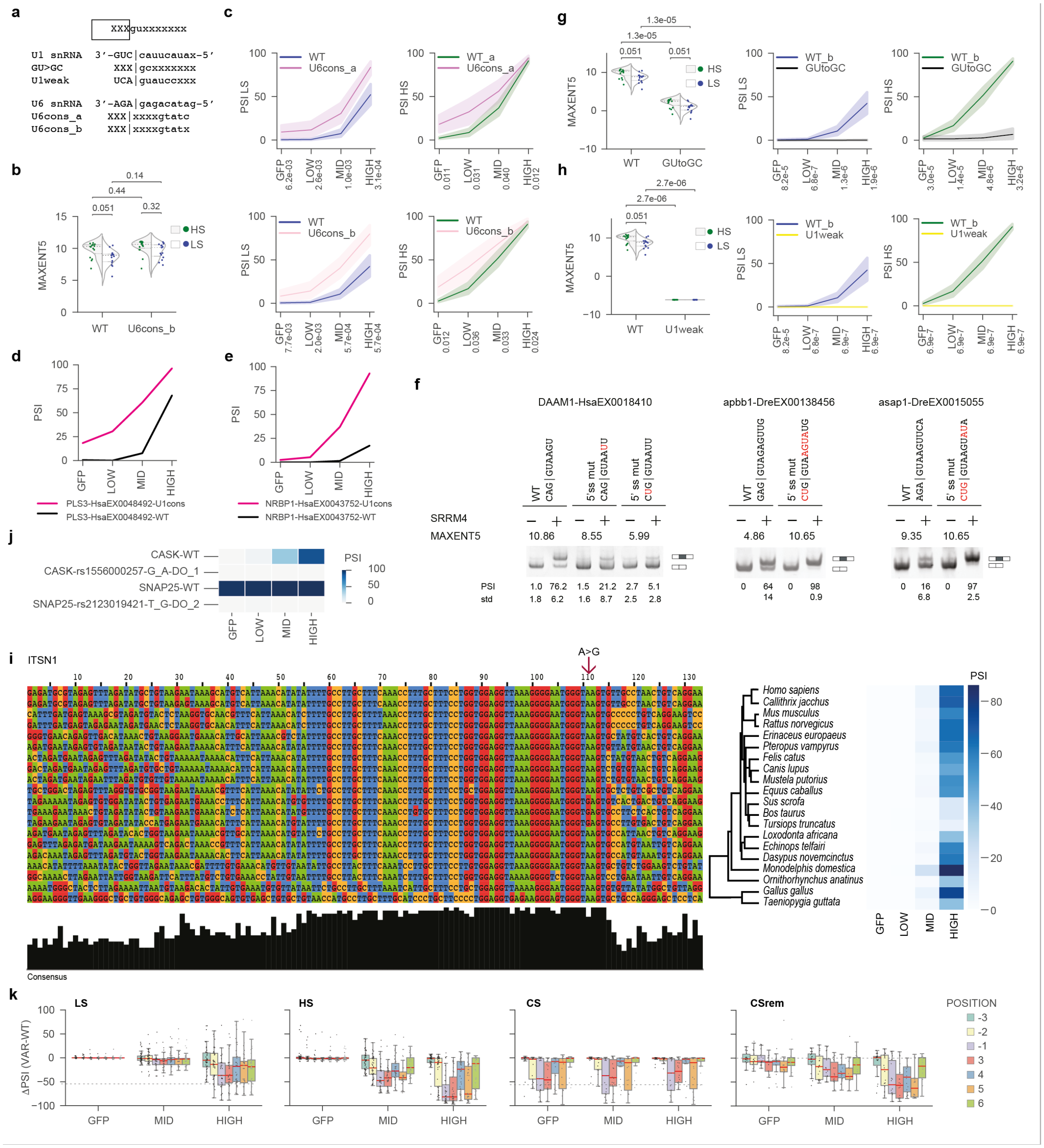

**Supplementary Fig. 6: The strength of the 5′ss contributes to splicing and sensitivity to SRRM4.** (**a**) Scheme of the mutations introduced to decrease complementarity to U1 snRNA or increase complementary to U6 snRNA. The top square represents the exon and its last three nts, the line represents the intron with the splice site “gu” and the following six nts. The sequence of U1 snRNA is provided as well the mutations performed to generate the variants named GT>GC (whereby position +2 is the intron is replaced from U to C) and U1weak (whereby 9 nts involved in base pairing to U1 snRNA have been replaced by the weak 5′ss associated with exon HsaEX6033277). In addition, positions +5 to +9 (U6cons_a) or +5 to +8 (U6cons_b) were mutated to enhance complementarity to U6 snRNA (whose sequence is provided above). (**b)** The lines within the violin represent the interquartile range of the maximal entropy score for 5′ss (MAXENT5) for the LS and HS events for the WT constructs and their corresponding U6cons_b variants. The median is represented by the dashed line, each dot represents an event. Statistics: Mann-Whitney-Wilcoxon test two-sided with Bonferroni correction. (**c**) The lines represent the PSI across four experimental conditions (GFP or LOW, MID, HIGH expression of SRRM4) for LS (left) or HS (right) events that are either WT or harbor mutations enhancing complementarity to U6 snRNA (U6cons_a, top panel or U6cons_b, bottom panel). Statistics: Mann-Whitney-Wilcoxon test. (**d,e**) The lines depict the PSI across four experimental conditions (GFP vs LOW, MID, HIGH expression of SRRM4) for PLS3-HsaEX0048492 (d) and NRBP1-HsaEX0043752 (e). The nature of the variant (WT or extended U1cons (U1cons_a)) is indicated by the color. The corresponding RT-PCR assays are provided in Fig. 6e. (**f**) RT-PCR assays showing the splicing patterns of different minigenes under control condition (-) or under expression of human SRRM4 (+) in HEK 293 cells. Each minigene was generated using the sequences from the endogenous events (upstream/downstream exons and introns). The variant name and the corresponding sequence of its 5′ss is indicated on top of the gel (3 last exonic nucleotide | 6 first intronic nucleotide). The mutations were engineered to either increase or decrease the complementarity to U1 snRNA and the corresponding maximal entropy score for 5′ss is reported (MAXENT5). The results correspond to a single replicate of the experiment. Each condition of DAAM1-HsaEX0018410 and apbb1-DreEX00138456/HsaEX0005055 was conducted in three biological replicates, while asap1-DreEX0015055/HsaEX0006155 was performed in four biological replicates, for which the means and standard deviations (std) are provided below each lane. (**g,h**) On the left side, the lines within the violin represent the interquartile range of the maximal entropy score for 5′ss (MAXENT5) for the LS and HS events for the WT constructs and their corresponding GU>GC (g) or U1weak (h) variants. Each dot represents an event. Statistics: Mann-Whitney-Wilcoxon test two-sided with Bonferroni correction. On the right side, the lines represent the PSI across four experimental conditions (GFP or LOW, MID, HIGH expression of SRRM4) for LS (left) or HS (right) events for WT and GU > GC (g) or WT and U1weak (h) variants. Statistics: Mann-Whitney-Wilcoxon test. (**i**) Alignment of ITSN1-HsaEX0032608 in 20 species spanning from primates to sauropsids for the last 93 nucleotides of the upstream intron, the microexon and 25 nucleotides of the downstream intron (as cloned in the context of the library). The level of conservation is depicted at the bottom with the black histogram and the nucleotides are counted from 1 to 133 at the top. The PSI values quantified across the four experimental conditions (GPP and LOW, MID, HIGH expression of SRRM4) are provided in the heatmap on the right side. The mutation from A to G in *sus_scrofa*, *bos_taurus* and *tursiops_truncatus* is indicated by the arrow. (**j**) The heatmap represents the effect of “likely pathogenic SNP” (ClinVar) mutations in four experimental conditions (GFP and LOW, MID, HIGH expression of

SRRM4) in the 5′ ss of the CASK-HsaEX0012545 HS event at position 1 (rs1556000257 G>A) and in the 5′ss of the CS event SNAP25-HsaEX3077184 at position 2 (rs2123019421 T>G). (**k**) Distribution of ΔPSI (VAR-WT) values between variants and their corresponding WT in GFP vs MID, HIGH expression of SRRM4 for LS, HS, CS and their corresponding shortened version events. Variants include deep mutagenesis of sequences involved in base pairing to U1 snRNA including positions -3, -2, -1 at the 3′ end of the exon and positions 3 to 6 of the intron.

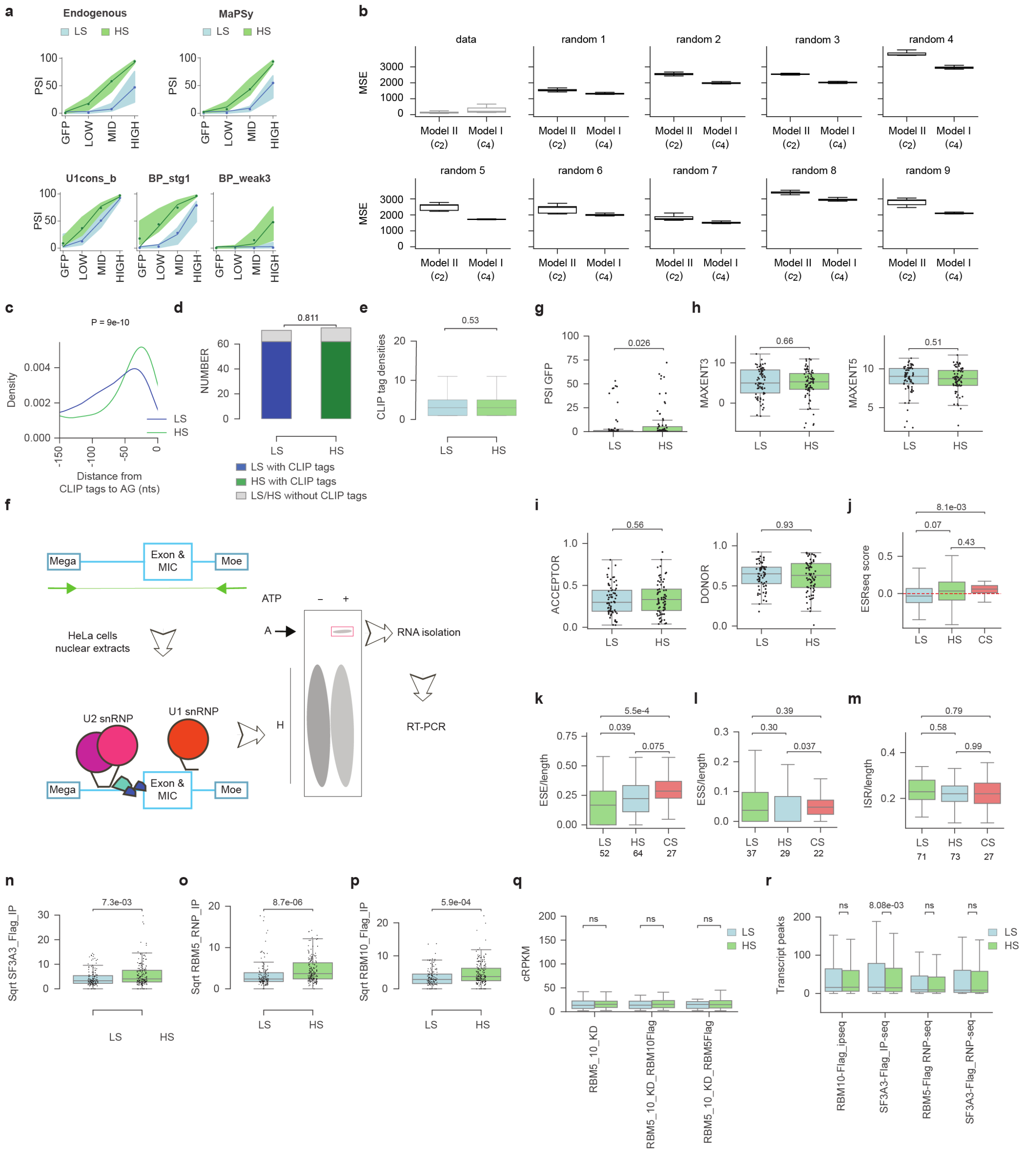


**Supplementary Fig. 7: Basal spliceosomal recruitment *in vitro* and *ex-vivo.***

(**a**) Fit of Model I to the data. Note that the model cannot explain the differences in the GFP condition among sequence groups. The model is the line, the dots the median of the data, and the shaded area the interquartile range. Same fit procedure as in Fig. 7c-e, but assuming that groups differ in $c_4$ instead of $c_2$. (**b**) Distribution of MSE (Mean Squared Error) for the best parameter sets found in each of 500 independent optimization runs, assuming groups differ in either $c_2$ or $c_4$, when the model was fitted either to the experimental data or to 9 random datasets with synthetic data constructed by permuting the median experimental PSI values across experimental conditions. (**c**) Frequency of the distance between the SRRM4 CLIP tags and the dinucleotide AG. Statistics: Mann-Whitney-Wilcoxon test. (**d**) Number of events with or without SRRM4 CLIP tags in the 200 nts preceding the microexons. Statistics: two-sided Fisher's Exact test. (**e**) SRRM4 CLIP tag densities in the last 150 nts of the upstream intron of endogenous HS and LS events, data from[57]. Statistics: Mann-Whitney-Wilcoxon. (**f**) Representation of the experimental approach used to investigate the efficiency of A spliceosomal complex formation *in vitro*. Briefly, RNA corresponding to the sequences cloned in Fig. 1c were transcribed and incubated in HeLa cells nuclear extracts under splicing conditions[10]. The H and A spliceosomal complexes were resolved by electrophoresis and RNA contained in A complex was isolated. The bands cut from the gels are highlighted by the red squares. Both the input and isolated RNA population (output) were amplified by RT-PCR assays and analyzed by deep-sequencing. Primers used for both input and output amplifications are depicted with the green arrows and amplicons in green line. (**g-i**): For 71 LS and 73 HS events: (g) PSI in GFP condition, (h) MAXENT3, MAXENT5 entropy scores and (i) SpliceAI scores of acceptor and donor sites. (**j-m**) For wild-type sequences of 71 LS microexons, 73 HS microexons and 27 42 nts long CS exons Mean ESRseq score (j), Number of ESE (k) or ESS (l) elements normalized by the exonic length and of ISR (m) elements identified in the 93 last nucleotides of the upstream intron and 25 first nucleotides of the downstream intron normalized by the total intronic length. Statistics: Mann-Whitney-Wilcoxon. Fisher tests for contingency tables: For ESE: LS-vs-HS, P = 0.035; LS-vs-CS= 1.3e-03; HS-vs-CS = 0.108. For ESS: LS-vs-HS, P = 0.181; LS-vs-CS= 0.010; HS-vs-CS = 2.5e-4. (**n-p**) The boxes represent the square root of the peak intensities of U2 snRNP binding in the 93 nts preceding the LS and HS microexons (Fig. 7g) as a result of either SF3A3_Flag_IP-seq (n), RBM5_RNP-seq (o) or RBM10_Flag_IP-seq (p). Statistics: Mann-Whitney-Wilcoxon test. (**q**) Expression of genes (in cRPKMs) presented in (Fig. 7f,g, and Supplementary Fig. 7g-h). (**r)** U2 snRNP peaks detected at the full transcript level. Statistics for f and g: Mann-Whitney-Wilcoxon test two-sided with Bonferroni correction (ns: non significant).

### SUPPLEMENTARY METHODS

The final pools of plasmids bearing the variants have the following structure: pt1 BC (34-nt long) *exon 1* intron 1 with Mega **VARIANT** (*93 nt- EXON - 25 nt*) Moe intron 2 *exon 3* 25 nts downstream intron pt2. The corresponding sequence is:

`gtcgacgacacttgctcaacNNNNAGCTNNNNTCAGNNNNTAGCNNNCAGTNNN`

*`GAATGTCTACAAGGATTATCGGCAGCTGGAGCTAGCCTGTGAGACACAGGAGGAGGTGGACAGCTGGAAGGCCT`*

*`CCTTCCTGAGGGCTGGCGTGTACCCTGAGCGTGTTGGG`*

GTGAGTGGCAGGGCAAGGAGAGGAAGGGCAAGCATGATCCTAGGGCCCCTGGGGCACCATCCTCAGTGATGCCA
AGTCATGCCATGTTTCCGAGCCCTTATTTGGCTCAGAAATAATAGGAATCCTCCCCCCTACCCACTCTGGGGGT
GGGAACAGAGATAAGTCTCCTGGTATTCCCATCCTTCTCCAGTGGCAGTTTTGTGTCTCTGTCTCTGTTGCAGA
TGGCATTTCCTCCATCCCCTTTCTATGATGGTAGTTTCTTGTGACTGCCTTCTCTTTTCTCCCCTATTTCACTG
TGGCGATGTCTAGAGTAGCCTGAGAGTTGATGGGATAAGACGGTAGGC---**VARIANT**---
TCGTAGCACGTCACGGTTGGAGCTCCAGCCAGGTTTTCAAGCAAGGGACCTGGAGATGTTCTTTTCTAATTTCT
GGATTGGGGCCAGGCGCAGTGGCTCACACCTGTAAACCCAACACTTTGGGAGGCCGAGGTGGGCGGATCACAAG
GTCAGGAGTTCGAGACCAGCCTGGCCAACACGGTGAAACCCCACCTCTACTAAAAATACAAAAATTAGCCAGGC
ATGGTGGTGCGCGCCTGTAATCCCAGCTACTCAGGAGGCTGAGGCAGGAGAATCGCTTGAACCCAGGAGGAGGT
TGCAATGAGCCAATACAGCACCACTGCACTCCAGCCTGGGTGACAAAGCAAGACTTTCTCAAAATAATAATAAT
AATAATTTCTGGATTGGGAAATTGAGGCAAATTCTAGGCACTAGAGTCAGAACCAAGACAAGGCTGAATCAGGG
GAGTCTAGGGTCCTGAGAGGCACAGGGATGCGGAGCCAGGTATGTATTCAGGCCAGTCGCTTCTCTCTGTGCCT
CAATATTCTGGGTACCCTTGGAGGGGCTGAGATCCTGGGGATGCCTGGAGCCTGGCTGCATGGCCTGGCCACCT
GATGCCCTTGTGTTCTCCATGGCAG
*GCCAGCGAGACCGAGGAGAATGGCTCCGACAGC**c**TCATGCATTCCATGGACCCACAGCTGGAACGGCAAGTGGA*
*GACCATCCGGAATCTTGTGGACTCATACATGGCCATTGTCAACAAGACCGTGAGGGACCTCATGCCCAAGACCA*
*TCATGCACCTCATGATTAACAAT*
GTGCGTGCTCCACTGCATGGGGGCActactgattcgatgcaagc

### SUPPLEMENTARY TABLES

All tables are provided as standalone excel documents.

**Supplementary Table 1:** related to Fig. 1 and Supplementary Fig. 1. List of the 234 events (EventID from https://vastdb.crg.eu/) selected in the first library (T0) and their membership as lowly sensitive (LS), highly sensitive (HS), constitutively spliced (CS), cryptic (CR) or control exons.

**Supplementary Table 2:** List of the 43 events selected for libraries T1, T2, T3, T4 and their membership as LS, HS, CS and CR. Yes/No indicates whether the events were considered in the respective library including T0. HsaEX0002213, present in each library, was discarded from the analyses.

**Supplementary Table 3:** List of species from which sequences of microexons and surrounding intronic sequences were retrieved, together with the time of split from the human lineage in millions of years ago (MYA), acronyms used in the variant names, and the corresponding nodes (Fig. 1e, Supplementary Fig. 1c,e).

**Supplementary Table 4:** List of the 36-nt-long sequences used to elongate microexons. The names, sequences, alternative names, resources used for the design and libraries in which each sequence was used are provided. ESRseq scores[44] were used to design sequences with the lowest possible score (Supplementary Fig. 2a,b). In addition, sequences bearing AG, GT, TGC and tandem sequence repeats of 3 or more nucleotides were discarded.

**Supplementary Table 5:** Description of the type of mutations performed for variant generation. Classification is based on the location where the mutation is done in the pre-mRNA, either at the 3′ss (UPINT), the exon (VAR_SEQ) or the 5′ss (DOINT). Combinations of mutations in different parts of the pre-mRNA are also reported. Extension in VARIANT name and description of the mutation is provided for each type.

**Supplementary Table 6:** Results of one sample t-tests for Fig. 3b,c and 5b-e.

**Supplementary Table 7:** Exonic Splicing Regulators (ESR) and Branch Point (BP) sequences used for variant generation. SVM_BP scores are included for each BP sequence.

**Supplementary Table 8:** A/H ratio of A complex formation on ITSN1 for five biological replicates (related to Fig. 6f).

**Supplementary Table 9:** Resources (softwares and websites) used to recover information related to splicing features, evolution and github resources (VastDB, *vast-tools*, maxent3, maxent5, svm-bp, github for MaPSy analyses in this study and mathematical model).

**Supplementary Table 10:** related to Fig. 1. PSI of endogenous events as provided by *vast-tools* under 9 experimental conditions (HeLa SRRM3 and HEK 293 SRRM4 from Fig. 1 and Supplementary Fig. 1) with the classification of LS (71 events), HS (73 events), NR/Others (33 events), CR (30 events), CS (27 events). Columns with "*-Q" contain quality information for each sample, as provided by *vast-tools* (https://github.com/vastgroup/vast-tools?tab=readme-ov-file#combine-output-format). Missing values are reported as NA. Samples used were generated in[4] (HeLa FLip IN cells expressing GFP or Srrm3); in[10] (HEK 293 Flp-In T-REx expressing GFP) and in this study (CL_HEK293_SRRM4_CONT, CL_HEK293_SRRM4_LOW, CL_HEK293_SRRM4_MID, CL_HEK293_SRRM4_HIGH).

**Supplementary Table 11:** List of the publicly available RNA-seq datasets used in this study.

**Supplementary Table 12:** List of the tissues from the different species represented in Supplementary Fig. 1d.

**Supplementary Table 13:** Table related to the selection of the events from MaPSy T0 to T1,2,3,4

**Supplementary Table 14:** It contains the information of all the variants used in each of the 5 libraries. The columns correspond to: EVENT, VARIANT, OLIGO (the oligo sequence is provided without 20 nts of Mega / 20 nts of Moe added for ordering to TWIST Bioscience, i.e., 5′ end with Mega: 5′-GATGGGATAAGACGGTAGGC-3′ and 3′ end with Moe: 5′-TCGTAGCACGTCACGGTTGG-3′, Fig. 1c), UPINT (93 nts of the upstream intron), VAR_SEQ (sequence of the exon), DOINT (25 nts of the downstream intron), VAR_LEN (exonic length), LME (Libraries' number the variant corresponds to). CS events bearing "-rem" in the variant names correspond to shortened versions of the original CS events.

**Supplementary Table 15:** Pairwise matching of variants with identical sequences in library T4, belonging to different mutation groups. This information was used to reassign PSI in the final PSI table based on PSI for VARIANT (reported in the first column).

**Supplementary Table 16:** For each library, total number of WT events, number of unique OLIGO sequences and median number of barcodes per VARIANT (calculated from the set of unique barcode-variant associations used to quantify the inclusion of each variant).

**Supplementary Table 17:** Primer name, sequence and their use in this study.

**Supplementary Table 18:** Total number of reads per sample deposited to GEO (GSE276143).

**Supplementary Table 19:** PSI of a given VARIANT from libraries T0, T1, T2, T3, T4 in four experimental conditions (GFP expression and LOW, MID, HIGH expression of SRRM4). Missing values are reported as NA.

**Supplementary Table 20:** PSI and standard deviation of the biological replicates for each individual minigene presented.

**Supplementary Table 21:** Enrichment score (ES) of A complex formation performed under control condition (CTR) for variants in libraries T1, T2. Missing values are reported as NA.
